# Regulated repression, and not activation, governs the cell fate promoter controlling yeast meiosis

**DOI:** 10.1101/2020.01.13.904912

**Authors:** Janis Tam, Folkert J. van Werven

## Abstract

Intrinsic signals and cues from the external environment drive cell fate decisions. In budding yeast, the decision to enter meiosis is controlled by nutrient and mating-type signals that regulate expression of the master transcription factor for meiotic entry, *IME1*. How nutrient signals control *IME1* expression remains poorly understood. Here we show that *IME1* transcription is regulated by multiple sequence-specific transcription factors that mediate association of Tup1-Cyc8 co-repressor to its promoter. We find that at least eight transcription factors bind the *IME1* promoter when nutrients are ample. Remarkably, association of these transcription factors is highly regulated by different nutrient cues. Mutant cells lacking three transcription factors (Sok2/Phd1/Yap6) displayed reduced Tup1-Cyc8 association, increased *IME1* expression and earlier onset of meiosis. Our data demonstrate that the promoter of a master regulator is primed for rapid activation while repression by multiple transcription factors mediating Tup1-Cyc8 recruitment dictates the fate decision to enter meiosis.

## Introduction

The choice of whether to differentiate into another cell type is directed by multiple cell intrinsic and extrinsic environmental factors. These cues signal to master regulatory genes, which in turn control the initiation of cell differentiation programs. As a result, multiple signals are transformed into a binary decision: whether or not to undergo cell differentiation. How signalling cues coordinate a cell fate outcome has important implications for the understanding of development and diseases such as cancer.

Diploid budding yeast cells undergo an irreversible differentiation program called gametogenesis or sporulation during which a diploid cell gives rise to four haploid spores. The yeast gametogenesis program is characterized by one round of DNA replication and recombination, two consecutive chromatin segregation events called meiosis followed by spore formation ^1^. As a result, an ascus with four haploid spores is produced. In yeast, the decision to enter meiosis is controlled by a master regulatory transcription factor named <u>in</u>ducer of <u>me</u>iosis 1, *IME1* ^2, 3^. In the absence of *IME1*, cells cannot enter meiosis and produce gametes. Thus, understanding how *IME1* is regulated is key to understanding how the decision to enter meiosis or gametogenesis is made.

Transcriptional control mechanisms regulate *IME1* expression. The *IME1* gene has an unusually large promoter (over 2.2 kilobases) that integrates multiple signals to control *IME1* expression ^4^. Nutrient and mating type signals ensure that *IME1* is only expressed in the appropriate nutrient environment and in the correct cell type. *IME1* is expressed in cells harbouring opposite mating-type loci (*MAT*a and *MAT*α) ^5, 6^. In cells with a single mating type (*MAT*a or *MAT*α), the transcription factor Rme1 is expressed and induces transcription of the long noncoding RNA (lncRNA) *IRT1*, which in turn transcribes through the *IME1* promoter and thereby represses *IME1* expression ^7^. In *MAT*a/α diploid cells, a second lncRNA upstream of *IRT1* named *IRT2* interferes with *IRT1* transcription forming a positive feedback loop by which Ime1 promotes its own expression ^8^.

In addition to mating-type control, environmental cues also play a critical role in regulating *IME1* expression. In order to induce *IME1* transcription, diploid cells must be starved for glucose and nitrogen, and cells need to be respiring ^4, 9^. The glucose and nitrogen signals integrate at the *IME1* promoter. Several sequence elements in the *IME1* promoter are important for control of *IME1* transcription. For example, a distinct sequence element mediates *IME1* repression by glucose signalling, while other parts of the promoter respond to nitrogen availability ^10^. Notably, the transcription factor Sok2 controls *IME1* promoter activity via the glucose responding element ^11^. In addition, multiple other transcription factors contribute to regulation of *IME1* transcription ^12–14^. Over 50 transcription factors have a conserved consensus site in the *IME1* promoter ^12^. A genome-wide reporter screen revealed that about 30 transcription factors may directly or indirectly control *IME1* transcription ^12^. How different transcription factors and functional elements of the *IME1* promoter interact to control *IME1* expression and thus regulate the decision to enter meiosis is not well understood.

The nutrient control of *IME1* expression is mediated by multiple signalling pathways including PKA, TOR complex 1 (TORC1), AMP-activated protein kinase (AMPK) and mitogen-activated protein kinase (MAPK) ^15–17^. Inhibiting two signalling pathways, PKA and TORC1, is sufficient to induce *IME1* expression in cells exposed to a nutrient rich environment where *IME1* expression is normally repressed ^16^. Thus PKA and TORC1 signalling is essential for controlling *IME1* expression and hence the decision to enter meiosis (Figure 1A). Previously, we showed that Tup1 represses the *IME1* promoter under nutrient rich conditions. Tup1 is part of the Tup1-Cyc8 co-repressor complex, which is involved in repression of more than 300 gene promoters in yeast ^18–20^. During starvation, however, Tup1 disassociates from the *IME1* promoter and *IME1* transcription is concomitantly induced. Tup1 binding to the *IME1* promoter is controlled by PKA and TORC1 ^16^. When both signalling pathways are inhibited Tup1 dissociates from the *IME1* promoter. Thus understanding how Tup1-Cyc8 binds to the *IME1* promoter may reveal how nutrient signalling controls *IME1* expression and consequently how cells make the decision to enter meiosis.

**Figure 1.**
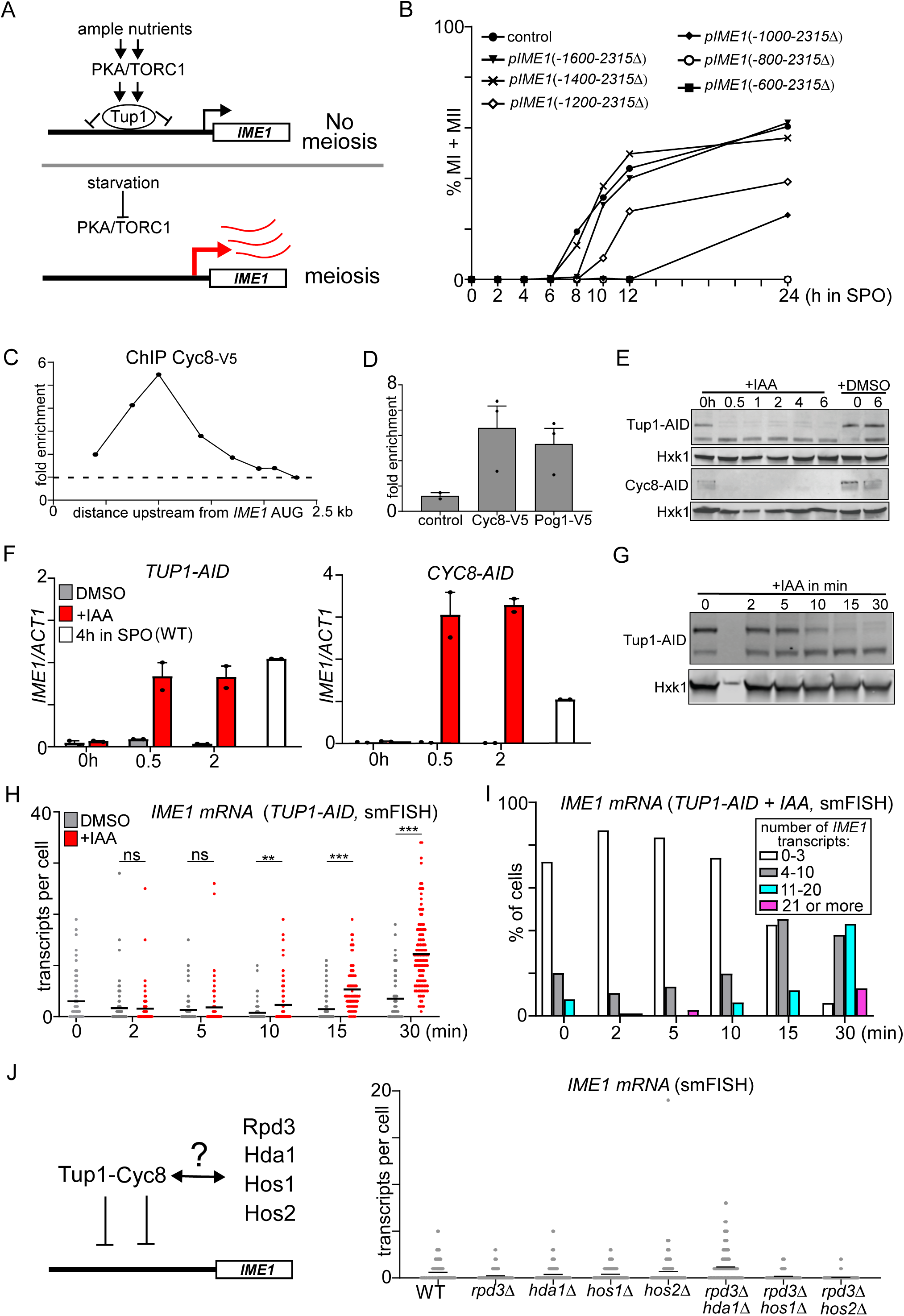
Tup1-Cyc8 repressor complex prevents activation of the *IME1* promoter independent of HDACs. (A) PKA and TORC1 integrate at the *IME1* promoter to control the association of the Tup1. Tup1, in turn, controls *IME1* repression. (B) The effect of truncations in the *IME1* promoter on the onset of meiosis. For the analyses we used diploid cells harbouring one copy of *IME1* deleted (control, FW4128), while different promoter deletion mutants were generated at the wild-type *IME1* locus (*pIME1*(−1600-2315Δ), FW3946; *pIME1*(−1400-2315Δ), FW3947; *pIME1*(−1200-2315Δ), FW3948; *pIME1*(−1000-2315Δ), FW3949; *pIME1*(−800-2315Δ), FW3950; *pIME1*(−600-2315Δ), FW3951). Control and mutant cells were grown to saturation in rich medium (YPD), grown for an additional 16 hours in pre-sporulation medium (SPO), and subsequently cells were shifted to sporulation medium. Samples were taken at the indicated time points, fixed, and DAPI masses were counted for the indicated time points to determine the percentage of cells that underwent meiosis (MI+MII). Cell harbouring two, three or four DAPI masses were classified as meiosis. (C) Cyc8, the interacting partner of Tup1, associates with the *IME1* promoter in the same region as Tup1. Diploid cells harbouring Cyc8 tagged with the V5 epitope (FW6381) were grown in rich medium (YPD) to the exponential phase. Cells were cross-linked, and chromatin extracts were prepared. Subsequently, Cyc8 bound DNA fragments were isolated by immunoprecipitation, reverse cross-linked, and quantified by qPCR. Eight different primer pairs were used to determine where Cyc8 binds in the *IME1* promoter. The signals were normalized over the silent mating locus *HMR*, which was not bound by Cyc8. (D) Similar as C except that the binding of Cyc8 (Cyc8-V5, FW6381), Pog1 (Pog1-V5, FW968), and wild type (control, FW1511) cells are displayed. The region around −1000 bp upstream of the AUG start codon of *IME1* was analysed for Cyc8 and Pog1 binding. The mean signals and SEM are displayed. (E) Diploid cells harbouring Tup1 or Cyc8 fused to auxin induced degron (AID) (*TUP1-AID*, FW5057 and *CYC8-AID*, FW6371) were grown till the exponential growth phase. Subsequently, cells were treated with IAA or DMSO for 0, 30 or 60 minutes, and samples for fixed with trichloroacetic acid. Protein extracts were prepared to determine Tup1 and Cyc8 protein levels by Western blot. As a control, Hxk1 levels were determined with anti-Hxk1 antibodies. (F) Similar as E except that *IME1* mRNA expression was determined by qPCR of *IME1* in cells *TUP1-AID* and *CYC8-AID* treated with IAA or DMSO. RNA was isolated, reverse transcribed, and quantified by qPCR. Signals were normalized by *ACT1*. Mean and SEM are displayed. (G) *IME1* expression in single cells upon Tup1 depletion. Tup1 protein levels (*TUP1-AID*, FW5057) detected by Western blot as described in E, except that levels at 0, 5, 10, 15 and 30 min after IAA. (H) Distribution of *IME1* transcript levels in single cells (*TUP1-AID*) treated with IAA or DMSO as described in G. Single molecule RNA fluorescent in situ hybridization (smFISH) was used detect *IME1* expression. Cells were fixed, and hybridized with *IME1* (AF594) and *ACT1* (Cy5) probes. Only cells positive for *ACT1* were used for the analyses. The mean number of transcripts per cell for each time point (black line) is displayed. At least 50 cells were used for the analysis. Unpaired parametric two-tailed Welch’s t-test with 95% confidence was used. Non-significant (ns) and P-values (** = ≤ 0.01, *** = ≤ 0.001) are indicated. (I) Same data as in H, except that the single cells data for *IME1* were binned according to expression levels. (J) Scheme of Tup1-Cyc8 interacting with histone deacetylases (HDACs) (left). *IME1* expression in diploid cells harbouring *rpd3*Δ, *hda1*Δ, *hos1*Δ, or *hos2*Δ single deletion mutants (FW8102, FW8426, FW8430, and FW8103) or *rpd3*Δ*hda1*Δ, *rpd3*Δ*hos1*Δ and *rpd3*Δ*hos2*Δ double deletion mutants (FW8457, FW8428, and FW8171) in single cells as determined by smFISH (right). The mean number of transcripts per cell for each time point (black line) is displayed. At least 100 cells were used for the analysis.

Here we set out to investigate how the Tup1-Cyc8 co-repressor complex regulates *IME1* transcription. In short, we found that regulated repression by multiple sequence specific transcription factors mediating the association of Tup1-Cyc8 with the *IME1* promoter is the means by which *IME1* transcription is controlled. Our data further indicate that nutrient cues highly regulate the association of Tup1-Cyc8 interacting transcription factors with the *IME1* promoter, which is key to regulating *IME1* expression and thus the cell fate decision to enter meiosis in yeast. Our work provides a framework to understand how nutrient signals integrate at a cell fate promoter and thereby control a developmental decision in yeast.

## Results

To gain insight into how Tup1 association with the *IME1* promoter is controlled, we first examined whether the region of the promoter where Tup1 binds contains key regulatory elements for *IME1* transcription. Previously, we found that Tup1 associates between 800 and 1400 base pairs (bp) upstream of the *IME1* translation start site ^16^. If the region of the *IME1* promoter where Tup1 binds is also important for *IME1* activation, then deleting that part of the promoter should affect the onset of meiosis. We generated six truncation mutants with a 200 bp interval in the *IME1* promoter and examined the ability of these mutants to undergo meiosis (Figure 1B). The largest truncation mutant that underwent meiosis with comparable kinetics as wild-type cells harboured 1400 bp of the *IME1* promoter (*pIME1*(−1400-2315Δ)) indicating that this region harbours the regulatory elements required for complete activation of the *IME1* promoter (Figure 1B). In addition, we found that meiosis in *pIME1*(−800-2315Δ) was completely impaired, whereas *pIME1*(−1200-2315Δ) had a much milder effect on meiosis. The result suggests that a region between −800 and −1200 bp harbours regulatory elements essential for *IME1* promoter function. We also made smaller truncations in the promoter and found that the region between −800 and −850 bp contains regulatory elements important for *IME1* activation because a large fraction of *pIME1*(−850-2315Δ) cells underwent meiosis (Supplementary Figure 1). In conclusion, the region important for Tup1 binding to the *IME1* promoter is also required for transcription of *IME1*.

Tup1 forms a complex with Cyc8 ^19, 20^. The Tup1-Cyc8 co-repressor complex is conserved and plays various roles in regulating gene transcription ^21^. Like Tup1, Cyc8 has also been implicated in regulation of *IME1* expression ^22^. To investigate how Cyc8 regulates *IME1* expression, we first determined Cyc8 binding with the *IME1* promoter under nutrient rich conditions. We found that Cyc8 peaked in the same region as Tup1 in the *IME1* promoter (Figure 1C and 1D). These data indicate that Tup1 is in a complex with Cyc8 at the *IME1* promoter.

Various models for Tup1-Cyc8 mediated repression of target gene promoters have been described ^21, 23, 24^. It has been proposed that Tup1-Cyc8 primarily regulates promoters by masking transcription factors from recruiting co-activators ^25^. Thus, Tup1-Cyc8 interacts with transcription factors with activation potential at promoters. We hypothesized that if Tup1-Cyc8 represses the *IME1* promoter by shielding co-activators, then transcriptional activators should be at the promoter under repressive conditions. To test this, we measured under nutrient rich repressive conditions the association of a known transcriptional activator of *IME1*, Pog1 ^7^. We found that Pog1, like Tup1-Cyc8, is indeed enriched at the *IME1* promoter under repressive conditions (Figure 1D). We conclude that Tup1-Cyc8 complex associates with the *IME1* promoter, and possibly masks transcriptional activators such as Pog1.

Various transcription factors have been implicated in controlling in *IME1* expression^12^. To further examine whether transcriptional activators are readily present at the *IME1* promoter, we measured *IME1* expression after depletion of Tup1 or Cyc8. We reasoned that if Tup1-Cyc8 represses the *IME1* promoter by restraining activating transcription factors, then depletion of Tup1 or Cyc8 should allow activators present to concomitantly induce *IME1* transcription. We used the auxin inducible degron (AID) system (*TUP1-AID* and *CYC8-AID*) and treated cells with indole-3-acetic acid (IAA) to achieve rapid protein depletion in cells ^26^. Rapid and sustained depletion of Tup1 and Cyc8 was achieved within 30 minutes after IAA treatment (Figure 1E).

Strikingly, *IME1* transcript levels strongly increased concurrently, and were comparable to those in wild-type cells entering meiosis when *IME1* expression is typically at its peak (Figure 1F). These data show that Tup1-Cyc8 repressor complex is pivotal for repressing the *IME1* promoter under nutrient rich conditions. Our data further suggest that the default state of the *IME1* promoter is active due to the presence of transcriptional activators, which are restrained by Tup1-Cyc8 under repressive conditions.

Although the transcriptional activator Pog1 is already bound in repressive conditions the at *IME1* promoter, it is possible that other transcriptional activators factors associate with the *IME1* promoter after Tup1-Cyc8 dissociates. This may results in a delay between Tup1-Cyc8 depletion and activation of *IME1* transcription. We therefore decided to monitor *IME1* expression by single molecule RNA fluorescence in situ hybridization (smFISH). This technique can detect individual transcripts in single cells ^27^. We found that as soon as Tup1 was depleted, *IME1* transcripts were detected in single cells (Figure 1G, 1H, 1I, and Supplementary Figure 2A). An increase in *IME1* transcripts was detected 10 minutes after IAA treatment when Tup1 was partially depleted (Figure 1G). After 15 min of inducing Tup1 depletion, 5.3 *IME1* transcripts were detected per cell on average, and 12% of the cells (Tup1-AID+IAA) had more than 10 *IME1* transcripts compared to 2% in control cells (Figure 1H and 1I). At 30 minutes after IAA treatment, more than 55% of cells expressed more than 10 *IME1* transcripts. It is worth noting that the AID-tag fused to Tup1 had some effect on *IME1* expression in the absence of IAA as *IME1* transcript levels were increased by five-fold in *TUP1-AID* compared to wild-type cells (Figure 1H, 1I, and Supplementary Figure 2B and 2C). We conclude that there is little to no temporal delay between Tup1-Cyc8 depletion and *IME1* expression suggesting that transcriptional activators are bound or readily available for recruitment to the *IME1* promoter.

The Tup1-Cyc8 complex also controls gene promoters by altering chromatin states ^28–30^. Specifically, Tup1-Cyc8 directly interacts with class I and II histone deacetylases, which in turn confer repression through deacetylation of nucleosomes ^31–33^. For example, repression of the *FLO1* promoter is achieved by Tup1-Cyc8 mediated recruitment of Hda1 and Rpd3 ^34^. In *hda1*Δ *rpd3*Δ cells significant de-repression of the *FLO1* gene can be observed. To examine whether histone deacetylases mediate Tup1-Cyc8 repression at the *IME1* promoter, we generated single and double mutants of known Tup1-Cyc8 interacting histone deacetylases (Rpd3, Hda1, Hos1, and Hos2) and measured *IME1* expression levels by smFISH (Figure 1J and Supplementary Figure 2C). Deletion of individual HDACs (*rpd3*Δ, *hda1*Δ, *hos1*Δ, and *hos2*Δ) did not increase expression of *IME1*. About 10% of *rpd3*Δ*hda1*Δ cells expressed four or more *IME1* transcripts, a marginal increase when compared to Tup1 depleted cells (Supplementary Figure 2C). Two other double mutants (*rpd3*Δ*hos1*Δ and *rpd3*Δ*hos2*Δ) displayed no detectable increase in *IME1* expression. We conclude that HDACs that are known to interact with Tup1-Cyc8 play only a marginal role in repressing the *IME1* promoter.

The Tup1-Cyc8 interacts with DNA sequence specific transcription factors to form co-repressor complexes at promoters ^35–41^. These transcription factors facilitate Tup1-Cyc8 association with promoters and thereby mediate repression of target genes. Our observation that Tup1-Cyc8 is the key repressor of the *IME1* promoter prompted us to examine which transcription factors recruit Tup1-Cyc8 to the *IME1* promoter and how they control *IME1* transcription. First, we assembled a list of transcription factors previously reported to interact with either Tup1 or Cyc8. In addition, we examined the region of the *IME1* promoter (−600 to −1200 bp) where Tup1-Cyc8 binds, and scanned for sequence motifs among transcription factors known to interact with Tup1-Cyc8 (Figure 2A, Supplementary Figure 3A and 3B). We identified 13 candidate transcription factors that were known or implicated to interact with Tup1-Cyc8 and have binding sites in the *IME1* promoter (Figure 2A and Supplementary Figure 3A). We also included the transcription factor Sok2 in our analyses because it has been proposed to interact with Tup1-Cyc8 and Sok2 is known to directly repress *IME1* transcription ^11, 35^. After the curation of the list of transcription factors, we subsequently measured the binding of under nutrient rich conditions by epitope tagging each transcription factor and performing ChIP. Remarkably, eight transcription factors displayed enrichment at the *IME1* promoter (Figure 2B). As expected, a known regulator of the *IME1* promoter, Sok2, was strongly enriched ^11^. In addition, Phd1 (a paralog of Sok2) and Yap6 also displayed strong enrichment (Figure 2B). The transcription factor Sut1, which is known to interact with Cyc8, was also enriched ^38^. The transcription factors Mot3, Sko1, Nrg1, and Nrg2 displayed a milder enrichment, but their binding was above background levels. For all transcription factors that displayed enrichment, we also assessed their binding to other parts of the *IME1* promoter (Figure 2C). Transcription factors exclusively co-localised with Tup1-Cyc8 in the same region of the IME1 promoter around 1000 bp upstream of the *IME1* start codon. Thus, at least eight transcription factors that are known or have been implicated to interact with Tup1-Cyc8 associate with the *IME1* promoter.

**Figure 2.**
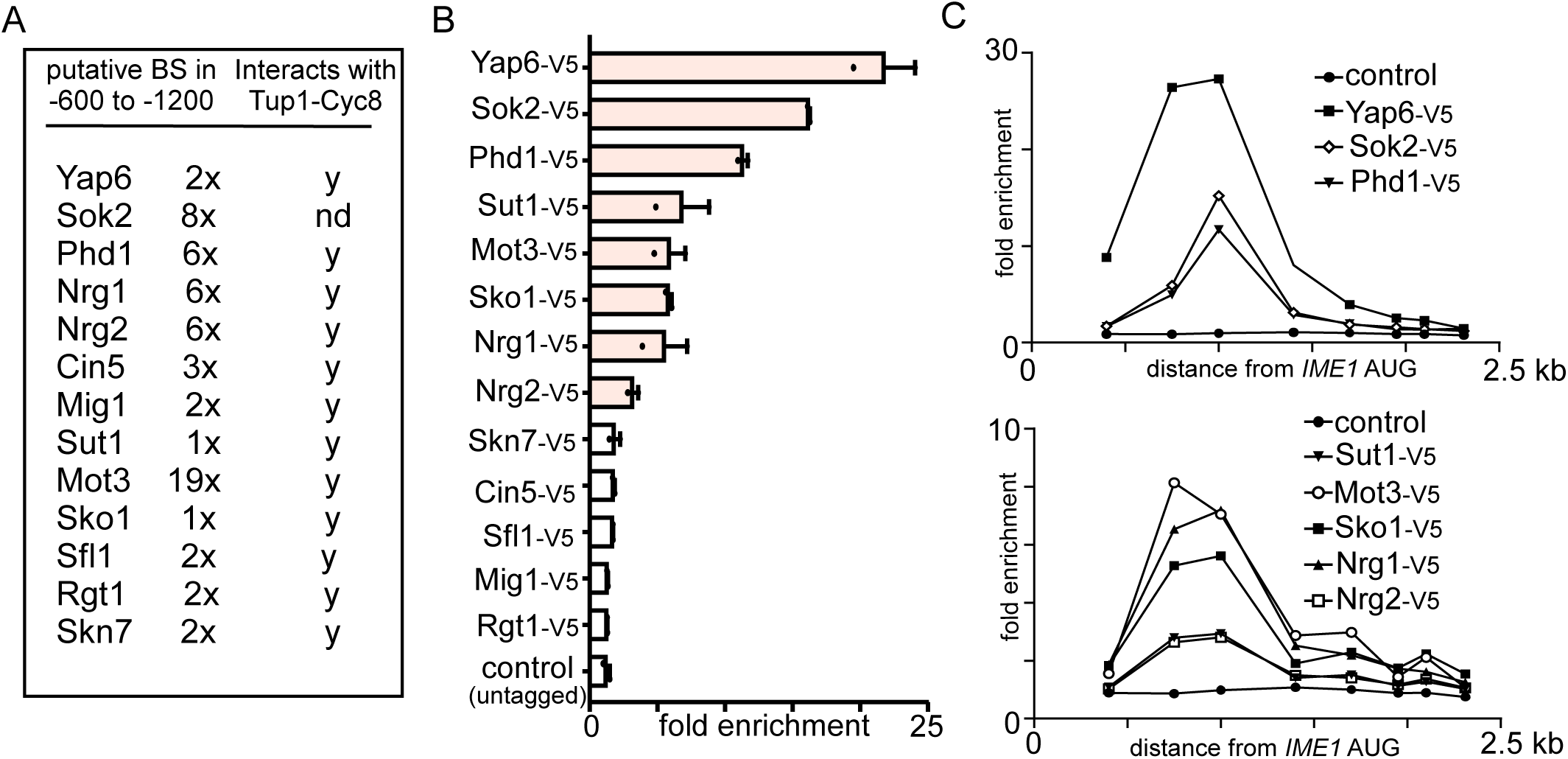
Multiple Tup1-Cyc8 interacting transcription factors associate with *IME1* promoter. (A) Table of transcription factors that have been implicated to interact with Tup1-Cyc8, and have motif sequences within the *IME1* promoter. (B) Multiple transcription factors associate with *IME1* promoter in the same region where Tup1-Cyc8 binds. Transcription factors listed in A were tagged with V5 epitope tag. Diploid cells haboring V5-tagged transcription factor (*YAP6-V5*, FW3833; *SOK2-V5*, FW5638; *PHD1-V5*, FW4466; *SUT1-V5*, FW6974; *MOT3-V5*, FW4383; *SKO1-V5*, FW4389; *NRG1-V5*, FW4393; *NRG2-V5*, FW4396; *SKN7-V5*, FW4399; *CIN5-V5*, FW7072; *SFL1-V5*, FW7070; *MIG1-V5*, FW4665; *RGT1-V5*, FW4386) and control cells (untagged, FW1511) were grown till exponential growth. The region around −1000 bp upstream AUG of *IME1* was analysed for binding by ChIP. The signals were normalized over *HMR*. Mean and SEM are displayed. (C) Similar as B, except that transcription factors that showed enrichment in B were analysed for binding throughout the whole *IME1* promoter. Eight primer pairs were used for the analyses.

Next, we examined whether the candidate transcription factors are responsible for recruiting Tup1 to the *IME1* promoter. We reasoned that candidate transcription factors should associate independently of Tup1-Cyc8 to the *IME1* promoter, while the binding of transcription factors and Tup1-Cyc8 should depend on the presence of specific sequence motifs (Figure 3A). First, we depleted Tup1 (*TUP1-AID*+IAA), and measured binding for a subset of the bound transcription factors (Supplementary Figure 4A). Except for Sko1, the binding of the transcription factors to the *IME1* promoter was not affected by Tup1 depletion (Figure 3B). Thus, multiple transcription factors known to interact with Tup1-Cyc8 associate with the *IME1* promoter independently of Tup1-Cyc8. Second, we examined whether the candidate transcription factors contribute to Tup1-Cyc8 recruitment. We mutated putative binding sites of the transcription factors that showed binding to the *IME1* promoter. To do so, we generated a construct containing the full-length promoter, followed by sfGFP and the *IME1* gene (*pIME1*-*WT*). Subsequently, we mutated 103 nucleotides distributed across a region of 400 bp in the *IME1* promoter (*pIME1*-*bs*Δ) (Figure 3C and Supplementary Figure 4B). We integrated the constructs in the *TRP1* locus in cells harbouring a deletion of the endogenous *IME1* gene and promoter sequence. Tup1 association with the *IME1* promoter was nearly completely lost in *pIME1-bs*Δ cells (Figure 3C). As expected, control cells (*pIME1-WT*) did not show a decrease in Tup1 binding. Finally, we assessed how *IME1* expression and the onset of meiosis is affected in *pIME1-bs*Δ cells. Surprisingly, *IME1* expression in *pIME1-bs*Δ cells was reduced, suggesting the regulatory elements essential for Tup1-Cyc8 recruitment are also important for *IME1* activation (Figure 3D). In conclusion, DNA sequence motifs of transcription factors bound to the *IME1* promoter are required for Tup1-Cyc8 association with the *IME1* promoter.

**Figure 3.**
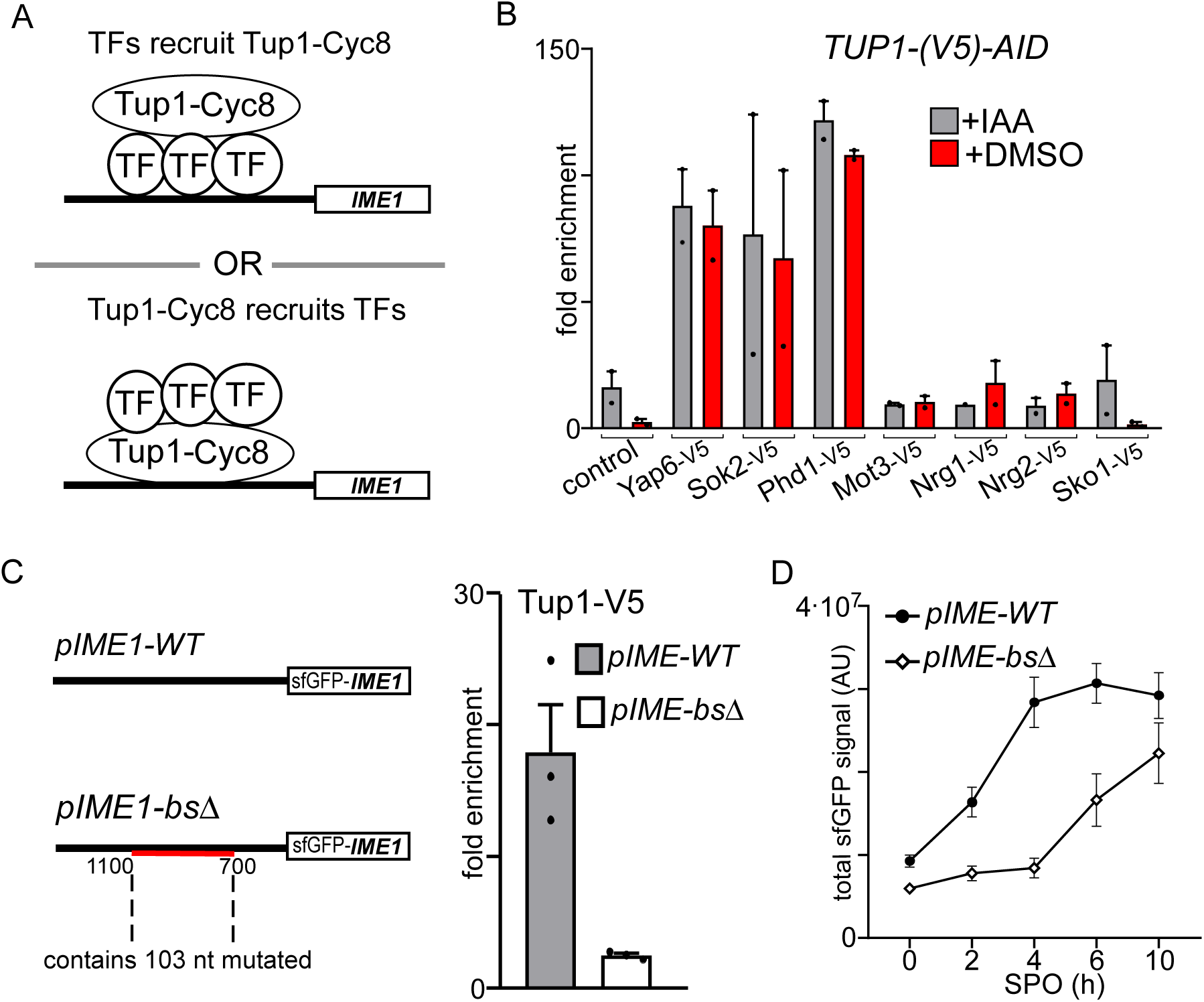
Tup1-Cyc8 is recruited by transcription factors associated with the *IME1* promoter. (A) Scheme displaying how transcription factors and Tup1-Cyc8 could interact at the *IME1* promoter. (B) Diploid cells harbouring *TUP1-AID* and V5-tagged transcription factors (*YAP6-V5*, FW4214; *SOK2-V5*, FW4218; *PHD1-V5*, FW5056; *MOT3-V5*, FW4229; *NRG1-V5*, FW4230; *NRG2-V5*, FW5055; *SKO1-V5*, FW4224) were grown to exponential phase. As a control, *TUP1-AID* cells were also included, which also harbours a V5 tag. Cells were either treated with IAA or DMSO, and the binding of each transcription factors was determined by ChIP. Mean signals and SEM are displayed. (C) Binding of Tup1 is affected in cells lacking transcription factor binding motifs in *IME1* promoter. Diploid cells with a chromosomal deletion of the *IME1* locus, and an integrated plasmid that contained the full *IME1* gene fused at amino terminus with sfGFP and promoter (*pIME1-WT,* FW5370) or the same construct with all the candidate motif sequences mutated *(pIME1-bs*Δ, FW5372) were used for the analysis. These cells also harboured *TUP1-V5*. The binding of Tup1 was determined by ChIP. Mean and SEM values are displayed. (D) Expression of Ime1 during entry in meiosis in *pIME1-WT* and *pIME1-bs*Δ. Strains described in C were grown till saturation in rich medium (YPD), grown for an additional 18 hours in pre-sporulation medium, and subsequently shifted in sporulation medium (SPO). The levels of Ime1 expression was determined by imaging and quantifying fluorescent signal generated by sfGFP-Ime1 in single cells. At least 50 number of cells were quantified. The mean signal and error bars representing the 95% confidence interval are displayed.

So far, our analyses of the *IME1* promoter showed that multiple transcription factors and multiple binding sites are essential for Tup1-Cyc8 recruitment. Next, we examined how the different transcription factors control *IME1* expression and mediate Tup1-Cyc8 recruitment under nutrient rich conditions. First, we assessed how the paralogs Sok2 and Phd1 control *IME1* expression. *sok2*Δ cells displayed a negligible increase in *IME1* expression (average transcripts per cell: 0.6 for *sok2*Δ versus 0.3 for control) (Figure 4A and 4B). In the *sok2*Δ*phd1*Δ double mutant, *IME1* expression was marginally increased (average transcripts per cell: 2.2) and about 5% of cells displayed more than 10 transcripts per cell suggesting that Sok2 and Phd1 play redundant roles in tightly repressing the *IME1* promoter (Figure 4A and 4B). We also measured *IME1* expression in *yap6*Δ cells or in combination with the *sok2*Δ*phd1*Δ mutation. *IME1* repression was not affected in cells containing *yap6*Δ, but in the *sok2*Δ*phd1*Δ*yap6*Δ triple deletion mutant *IME1* expression was increased (average transcripts per cell: 2.8 for *sok2*Δ*phd1*Δ*yap6*Δ versus 2.2 for *sok2*Δ*phd1*Δ) (Figure 4A and B). About 8% of *sok2*Δ*phd1*Δ*yap6*Δ cells expressed more than 10 *IME1* transcripts per cell. It is worth noting that *IME1* transcript levels in *sok2*Δ*phd1*Δ*yap6*Δ cells were much lower than in cells depleted for Tup1 (Figure 1F and 1H) - suggesting that additional transcription factors contribute to *IME1* repression.

**Figure 4.**
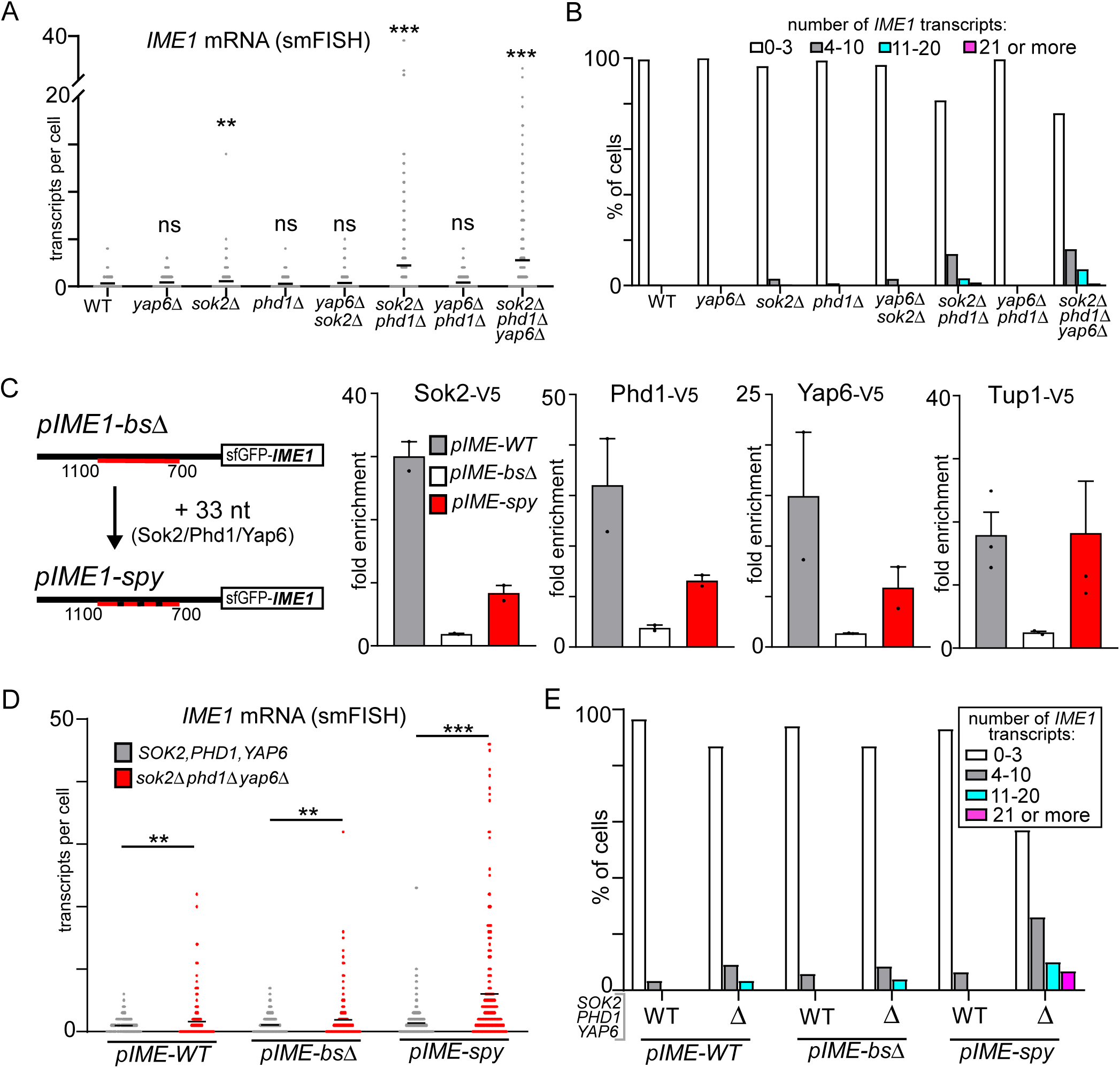
Multiple Tup1-Cyc8 interacting transcription factors repress *IME1* in nutrient rich conditions. (A) *IME1* expression in single cells harbouring: *sok2*Δ, *phd1*Δ or *yap6*Δ single mutants (FW3979, FW3991, FW3603); *sok2*Δ*phd1*Δ, *yap6*Δ*phd1*Δ and *sok2*Δyap6Δ double mutants (FW4710, FW4406, FW4239); *sok2*Δ*phd1*Δ*yap6*Δ triple mutant (FW4010). Cells were grown to exponential growth, were fixed, and hybridized with *IME1* (AF594) and *ACT1* (Cy5) probes. Only cells positive for *ACT1* were used for the analyses. The mean number of transcripts per cell for each time point (black line) is displayed. Between 190 and 300 cells were used for the analysis. Unpaired parametric two-tailed Welch’s t-test with 95% confidence was used. Non-significant (ns) and P-values (** = ≤ 0.01, *** = ≤ 0.001) are indicated. (B) Same data as in A, except that the single cells data for *IME1* were binned according to expression levels. (C) Sok2, Phd1, Yap6 binding in *pIME1-WT* (FW8081, FW8083, FW8079), *pIME1-bs*Δ(FW8087, FW8089, FW8085), and *pIME1-spy* (FW8093, FW8095, FW8091) as determined by ChIP. Mean and SEM values are displayed. (D) *IME1* expression in single cells as determined by smFISH in *pIME1-WT*, *pIME1-bs*Δ, and *pIME1-spy* in cells wild type for *SOK2*, *PHD1* and *YAP6* (FW5370, FW5372, FW7733) or harbouring the *sok2*Δ*phd1*Δ*yap6*Δ triple deletion mutant (FW7650, FW8420, FW8177). Unpaired parametric two-tailed Welch’s t-test with 95% confidence was used. P-values (** = ≤ 0.01, *** = ≤ 0.001) are indicated. (E) Same data as in D, except that the single cell data were binned according to *IME1* expression levels.

Our data demonstrate that Sok2, Phd1 and Yap6 associate with the *IME1* promoter and contribute to *IME1* repression in nutrient rich conditions. Yet, *IME1* expression was reduced in cells with DNA sequence motifs mutated (*pIME1-bs*Δ). One explanation is that the mutated binding sites in *pIME1-bs*Δ important for Tup1-Cyc8 recruitment also facilitate binding for transcriptional activators. Another possibility is that transcription factors important for Tup1-Cyc8 recruitment are also required for *IME1* activation. To discriminate between both possibilities, we generated a construct that contained binding sites for <u>S</u>ok2, <u>P</u>hd1, and <u>Y</u>ap6 (*pIME1-spy*), while the other transcription factor binding sites remained mutated (Figure 4C and Supplementary Figure 5). By combining *pIME1*-*spy* with *sok2*Δ*phd1*Δ*yap6*Δ, we could then determine whether Sok2, Phd1, and Yap6 are important for *IME1* activation or repression. Tup1 binding was restored in cells harbouring *pIME1-spy* (Figure 4C). Furthermore, Yap6, Sok2, and Phd1 were enriched at the *IME1* promoter in *pIME1-spy* cells but their binding was reduced compared to the wild-type promoter - suggesting that there are additional binding sites (Figure 4C). Next, we measured *IME1* expression in wild-type and *sok2*Δ*phd1*Δ*yap6*Δ mutant cells harbouring *pIME1-spy* (Figure 4D). We found that *IME1* expression was further de-repressed in cells harbouring both *pIME1-spy* and *sok2*Δ*phd1*Δ*yap6*Δ compared to cells expressing the three transcription factors (Figure 4D and 4E). About 17% of cells harbouring *pIME1-spy* and *sok2*Δ*phd1*Δ*yap6*Δ expressed more than 10 *IME1* transcripts per cell. As expected, the *sok2*Δ*phd1*Δ*yap6*Δ had only a mild effect on *IME1* levels in cells expressing *pIME1-WT* or *pIME1-bs*. We conclude that Sok2, Phd1 and Yap6 are important for repression of the *IME1* promoter, and play little role in *IME1* activation.

The Tup1-Cyc8 complex dissociates from the *IME1* promoter in cells exposed to nutrient starvation ^16^. We hypothesized that transcription factors interacting with Tup1-Cyc8 at the *IME1* promoter control Tup1-Cyc8 dissociation during *IME1* activation. For example, one possibility is that the physical interaction between Tup1-Cyc8 and transcription factors is altered during activation *IME1*. Alternatively, during activation of the *IME1* promoter transcription factors dissociate together with Tup1-Cyc8. To examine this, we measured the binding of the eight transcription factors during activation of the *IME1* promoter. In order to induce *IME1* expression and meiotic entry in a synchronous manner in all cells, we typically grow cells in rich medium conditions containing glucose until saturation, then shift and grow cells in pre-sporulation medium containing acetate. Cellular respiration is required for meiosis, and growth in medium with acetate but lacking glucose ensures that cells are respiring and not subject to repressive glucose signalling to the *IME1* promoter ^4, 9^. Subsequently, cells are starved by shifting them to sporulation medium (0.3% acetate), which induces *IME1* transcription and drives meiotic entry. First, we measured the binding of Tup1 and Pog1 at the *IME1* promoter prior to induction of *IME1* (0 hours in SPO), and during meiotic entry when *IME1* is induced (4 hours in SPO) (Figure 5A). Both Pog1 and Tup1 were enriched at 0 hours in SPO prior to *IME1* induction. As expected, during meiotic entry (4 hours in SPO) Tup1 dissociated from the *IME1* promoter completely while Pog1 binding was maintained albeit to a reduced level (Figure 5A). Next, we determined how binding of the different transcription factors with the *IME1* promoter was regulated. We found that all eight transcription factors, were enriched at the *IME1* promoter prior to induction of *IME1* (Figure 5A). Strikingly, five transcription factors showed no binding to the *IME1* promoter upon entry into meiosis, while three transcription factors (Yap6, Phd1, and Nrg1) displayed marginal enrichment (4 hours in SPO). These data suggest that Tup1-Cyc8 dissociates from the *IME1* promoter due to the loss of binding of multiple transcription factors. In conclusion, activation of *IME1* transcription correlates with the dissociation of transcription factors important for Tup1-Cyc8 recruitment, while a transcriptional activator remains present at the *IME1* promoter.

**Figure 5.**
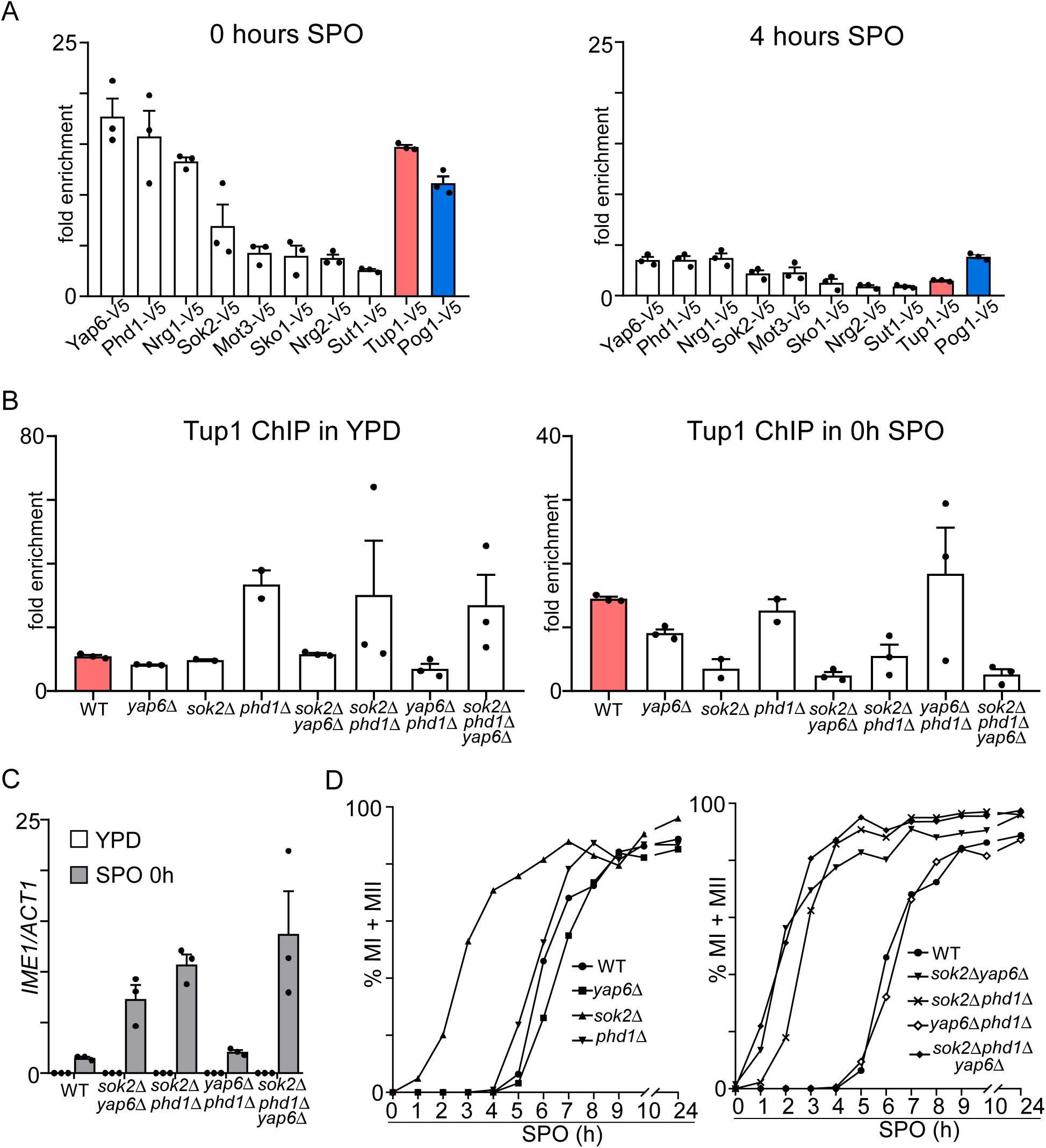
Sok2, Phd1 and Yap6 are important for repressing the *IME1* promoter prior entry into meiosis. (A) Transcription factors binding at the *IME1* promoter prior and during entry into meiosis. Diploid cells harbouring V5 tagged transcription factors (*YAP6-*V5, FW3833; *PHD1-V5*, FW4466; *NRG1-V5*, FW4393; *SOK2-V5*, FW5638; *MOT3-V5*, FW4383; *SKO1-V5*, FW4389; *NRG2-V5*, FW4396; *SUT1-V5*, FW6974; *TUP1-V5*, FW3456; *POG1-V5*, FW968) were used for the analyses. Samples for ChIP were taken at 0 and 4 hours in SPO. Mean and SEM values are displayed. (B) Tup1 binding at the *IME1* promoter at 0 hours in SPO (prior entry in meiosis) in wild-type cells or cells harbouring *sok2*Δ, *phd1*Δ or *yap6*Δ single deletion mutants (FW3979, FW3991, FW3603), and *sok2*Δ*phd1*Δ, *yap6*Δ*phd1*Δ and *sok2*Δyap6Δ double mutants (FW4710, FW4406, FW4239); *sok2*Δ*phd1*Δ*yap6*Δ triple mutant (FW4010). Mean and SEM values are displayed. (C) *IME1* expression in deletion mutants described in C. The signals were normalized over the *ACT1* gene. Mean and SEM values are displayed. (D) Meiosis in mutant strains described in B. Samples were taken at the indicated time points, fixed, and DAPI masses were counted for the indicated time points to determine the percentage of cells that underwent meiosis (MI+MII). Cell harbouring two, three or four DAPI masses were classified as meiosis. At least 200 cells were analysed.

We showed that Sok2, Phd1 and Yap6 were strongly enriched at the *IME1* promoter under nutrient rich conditions and prior to meiotic entry (Figure 2B, Figure 5A). In addition, in *sok2*Δ*phd1*Δ*yap6*Δ cells*, IME1* expression was partially de-repressed when nutrients were ample (Figure 4A and 4B). These observations indicate that Sok2, Phd1 and Yap6 are important transcription factors for *IME1* repression. Next, we investigated how the three transcription factors control Tup1-Cyc8 recruitment in different nutrient conditions. Specifically, we examined Tup1 binding to the *IME1* promoter in the absence of the three transcription factors in cells grown in rich medium containing glucose (YPD) and in acetate-containing medium prior to meiotic entry. We found that Tup1 binding to the *IME1* promoter was not affected in rich medium containing glucose in *sok2*, *phd1*, and *yap6* single/double/triple deletion mutants, which is in line with *IME1* expression data described in (Figure 5B, 4A, and 4B). This suggests that other transcription factors contribute to *IME1* repression via Tup1-Cyc8 when glucose is used by cells as the carbon source. In contrast, prior to meiotic entry (0h in SPO) Tup1 binding was significantly reduced in *sok2*Δ and *sok2*Δ*phd1*Δ cells, but not in *yap6*Δ and *phd1*Δ cells (Figure 5B). Strikingly, Tup1 association with the *IME1* promoter was reduced to nearly background levels in *sok2*Δ*yap6*Δ and *sok2*Δ*phd1*Δ*yap6*Δ cells at 0 hours in SPO. *IME1* expression inversely correlated with Tup1-Cyc8 recruitment to the *IME1* promoter (Figure 5C). For example, *sok2*Δ*yap6*Δ cells displayed significant de-repression of *IME1* expression prior to entry into meiosis compared to WT control cells. Likewise, cells harbouring *sok2*Δ*phd1*Δ or *sok2*Δ*phd1*Δ*yap6*Δ also displayed de-repression of *IME1* expression at 0 hours in SPO (Figure 5C). Finally, we examined how Sok2, Phd1, and Yap6 mediated Tup1-Cyc8 recruitment to *IME1* promoter is important for meiosis. In general, the onset of meiosis in the different mutants correlated well with *IME1* expression levels (Figure 5D). Cells harbouring the *sok2*Δ or *sok2*Δ*phd1*Δ underwent meiosis much faster than wild-type cells. For example after 3 hours in SPO, 50% of *sok2*Δ cells completed at least one meiotic division, while this took more than 6 hours in wild-type cells. There was little effect on the onset of meiosis in the *yap6*Δ or *yap6*Δ*phd1*Δ mutants. In *sok2*Δ*yap6*Δ and *sok2*Δ*phd1*Δ*yap6*Δ cells the kinetics of meiosis was slightly faster than *sok2*Δ*phd1*Δ cells. Approximately 50% of cells underwent meiotic divisions within 2 hours in SPO for *sok2*Δ*phd1*Δ*yap6*Δ cells compared to 3 hours for *sok2*Δ*phd1*Δ cells.

During meiotic entry (4 hours in SPO), Yap6, Phd1 and Nrg1 showed some enrichment at the *IME1* promoter suggesting that they could contribute to activation of *IME1* transcription. Therefore, we also analysed how the onset of meiosis is affected in *sok2*Δ*phd1*Δ*yap6*Δ*nrg1*Δ cells (Supplementary Figure 6). We found no difference in meiosis between *sok2*Δ*phd1*Δ*yap6*Δ*nrg1*Δ and *sok2*Δ*phd1*Δ*yap6*Δ cells suggesting that these transcription factors do not contribute to activation of *IME1* transcription in cells induced to enter meiosis synchronously. Taken together, we conclude that Sok2, Phd1, and Yap6 direct Tup1-Cyc8 association to the *IME1* promoter to ensure timely expression of *IME1* in cells grown in acetate containing medium. Our data suggest that the *IME1* promoter is regulated by multiple Tup1-Cyc8 co-repressor complexes.

How do nutrient signals regulate the association and dissociation of Tup1-Cyc8 recruiting transcription factors with the *IME1* promoter? While it is known that nutrients mediate PKA and TORC1 signalling pathways and thereby control *IME1* expression, little is known about the mechanisms that mediate PKA and TORC1 signalling at the *IME1* promoter ^4, 16^. Regulated localization or abundance of transcription factors are prevalent mechanisms for controlling gene expression via PKA and TORC1 ^42, 43^. Among the transcription factors that associate with the *IME1* promoter, Sok2 protein abundance is regulated by glucose signalling via the Ras/PKA pathway ^11, 44^. To gain insight into how Tup1-Cyc8 recruiting transcription factors (Sok2, Phd1, and Yap6) dissociate from the *IME1* promoter, we sought to investigate whether their abundance or localization is altered during entry into meiosis. First, we examined protein expression levels in exponentially grown cells (E), cells grown to saturation (S), and in cells just prior to and during meiotic entry (0h and 4h SPO). The levels of Sok2, Phd1 and Yap6 as well as Cyc8 were reduced in cells grown to saturation compared to exponentially growing cells (Figure 6A). However, in cells grown until saturation, Sok2, Phd1 and Yap6 were bound to the *IME1* promoter (Supplementary Figure 7). Interestingly, reduced Cyc8 levels mirrored the reduced binding of Cyc8 at the *IME1* promoter (Figure 6A and Supplementary Figure 7). Importantly, Sok2, Phd1 and Yap6 abundance was not altered in cells between the 0 and 4 hour time points (in SPO) when *IME1* expression is induced. These data indicate that cellular changes in Sok2, Phd1 and Yap6 abundance are unlikely to explain the dissociation of these transcription factors from the *IME1* promoter during entry into meiosis. To determine whether these transcription factors are evicted from the *IME1* promoter due to nuclear export, we determined the cellular localization for each transcription factor. We fused mNeongreen to Sok2, Phd1, Yap6, Tup1 and Cyc8 (Supplementary Figure 8A). As expected, Sok2, Phd1, and Yap6 were concentrated in the nucleus. Neither the nuclear-to-cytoplasmic ratios nor total protein abundance in the nucleus were significantly altered in cells prior to (0h in SPO) and during entry into meiosis (4h in SPO) (Figure 6B and Supplementary Figure 8B). Furthermore, the overall localization of Tup1 and Cyc8 was also not altered during meiotic entry (Figure 6B and Supplementary Figure 8B). We conclude that cellular changes in protein abundance or localization cannot solely account for the dissociation of Tup1-Cyc8 and transcription factors from the *IME1* promoter during entry in meiosis.

**Figure 6.**
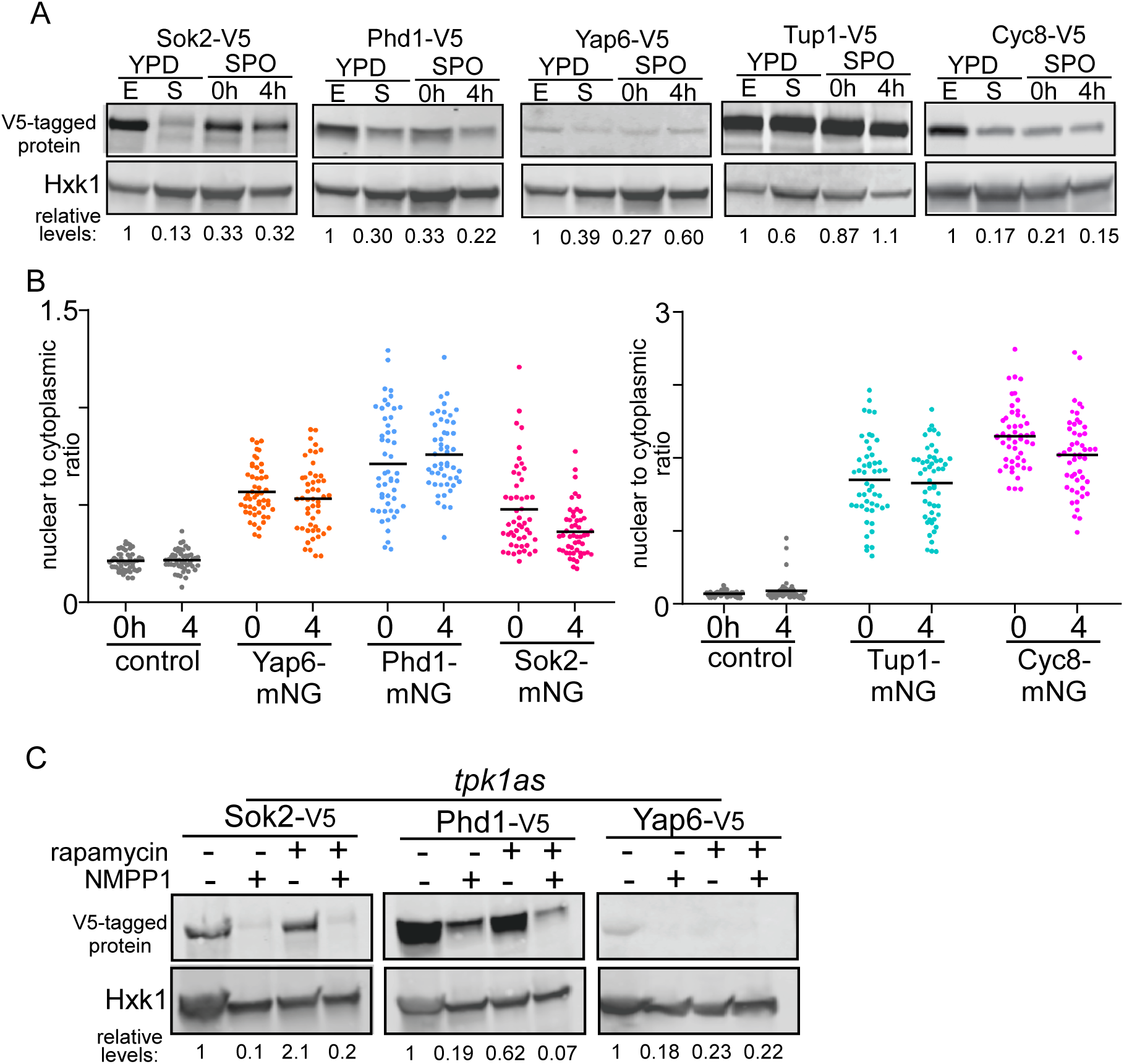
PKA and TORC1 control Sok2, Phd1 and Yap6 protein abundance. (A) Expression of Sok2, Phd1, Yap6, Tup1, and Cyc8 in cells grown to exponential phase or saturation in YPD, and 0 and 4 hours in SPO (*SOK2-V5*, FW5638; *PHD1-V5*, FW4466; *YAP6-V5*, FW3833; *TUP1-V5,* FW3456; *CYC8-V5,* FW6381). Samples for western blotting were taken at the indicated time points. Hxk1 was used as a loading control. Relative abundance is displayed for each transcription factor. (B) Nuclear to cytoplasmic ratio of Sok2-mNG (FW7475), Phd1-mNG (FW7477), Yap6-mNG (FW7473), Tup1-mNG (FW7644) and Cyc8-mNG (FW7642) prior (0 hours in SPO) and during entry into meiosis (4 hours in SPO). Each transcription factor was fused to mNeongreen (mNG). These cells also expressed a mCherry fused to SV40 nuclear localization signal (NLS) (mCherry-NLS). As a control, the signals of cells harbouring no mNG-tag (FW5199) are displayed. The black bar indicates the mean signal, and each point indicates a single cell measurement. (C) Similar as A, except that expression of Sok2, Phd1, and Yap6 was monitored after inhibiting PKA and TORC1. We used cells harbouring the *tka1-as* allele for inhibiting PKA activity with ATP analog 1NMPP1 (*YAP6-V5*, FW5453; *SOK2-V5*, FW5454; *PHD1-V5*, FW5528). TORC1 was inhibited with rapamycin. Cells were grown to exponential growth, treated with 1NMPP, rapamycin, or both for 4 hours. Sok2, Phd1 and Yap6 protein levels are normalized over Hxk1. Relative abundance with respect to YPD (E) is displayed.

There is evidence that signalling kinases can act locally at gene promoters. For example, Tpk1 and Tpk2 kinases of PKA can associate with specific gene promoters and regulate transcription locally ^45^. We reasoned that our measurements of transcription factor abundance and localization may be masked if PKA and TORC1 signalling is heterogeneous within cells. Therefore, perhaps inhibiting PKA and TORC1 altogether could reveal how both signalling pathways regulate transcription factors important for repressing the *IME1* promoter. To inhibit PKA we used the *tpk1-as* allele previously described, while we used rapamycin to inhibit TORC1 ^16^. Strikingly, Sok2, Phd1 and Yap6 protein levels were reduced between five to ten-fold when PKA was inhibited (*tpk1-as* + NMPP1) (Figure 6C). Inhibiting TORC1 activity with rapamycin also lowered Yap6 and Phd1, but not Sok2 levels. We conclude that inhibiting PKA and TORC1 affects the abundance of transcription factors important for repressing the *IME1* promoter, and coincides with Tup1 disassociation and activation of *IME1* transcription as described previously ^16^. Our data suggest that the association of transcription factors with the *IME1* promoter may be regulated locally by PKA and TORC1 signalling.

Our data indicate that at least eight transcription factors that are known or implicated to interact with Tup1 or Cyc8, associate with the *IME1* promoter (Figure 2B). Motif analyses revealed that there are 52 corresponding binding sites present in the region of the *IME1* promoter where Tup1-Cyc8 associates (Supplementary Figure 3). In addition, there are multiple binding motifs for almost every transcription factor present suggesting that multiple copies of each transcription factors bind to the *IME1* promoter (Supplementary Figure 3A). Why is there need for so many binding sites in the *IME1* promoter? One possibility is that the transcription factors are important for facilitating Tup1-Cyc8 recruitment under different nutrient conditions. With this logic, the repression of the *IME1* promoter can be maintained under various nutrient conditions, and will only be fully activated when all the nutrient signalling requirements are met. In agreement with this model, *IME1* expression was only marginally increased in *sok2*Δ *phd1*Δ*yap6*Δ cells grown in the presence of ample nutrients with glucose as the carbon source (YPD), while the *IME1* promoter was nearly completely de-repressed in *sok2*Δ*phd1*Δ*yap6*Δ cells grown in acetate-containing medium (Figure 4A, 4B, 5B and 5C). To examine how the different transcription factors respond to nutrient signalling at the *IME1* promoter more systematically, we measured their association under different nutrient conditions (Figure 7A). We grew cells until the pre-sporulation stage, and subsequently shifted cells to sporulation medium (SPO) (1), SPO plus 2% glucose (2), YP (3), and YP plus 2% glucose (YPD) (4). First, we measured Tup1 association with the *IME1* promoter. We found that in SPO plus glucose, Tup1 binding to the *IME1* promoter was partially restored (Figure 7B). The association of Tup1 with the *IME1* promoter was further increased in YP and was the highest in YPD growth medium. The transcriptional activator of *IME1*, Pog1, was enriched in all four nutrient conditions, but at higher levels in YP and YPD (Figure 7B). Interestingly, transcription factors important for Tup1-Cyc8 recruitment to *IME1* promoter responded to nutrient signals in distinct ways (Figure 7C). For example, Yap6, Sok2, Sko1 and Nrg1 did not associate with the *IME1* promoter in response to glucose, but their binding was restored due to nutrient cues present in YP. Phd1 binding partially recovered in the presence of glucose, and showed the strongest enrichment in cells exposed to YP and YPD. The carbon source, glucose, but not YP, induced association of Mot3 and Nrg2 with the *IME1* promoter, while Sut1 association with the *IME1* promoter was restored in YPD only.

**Figure 7.**
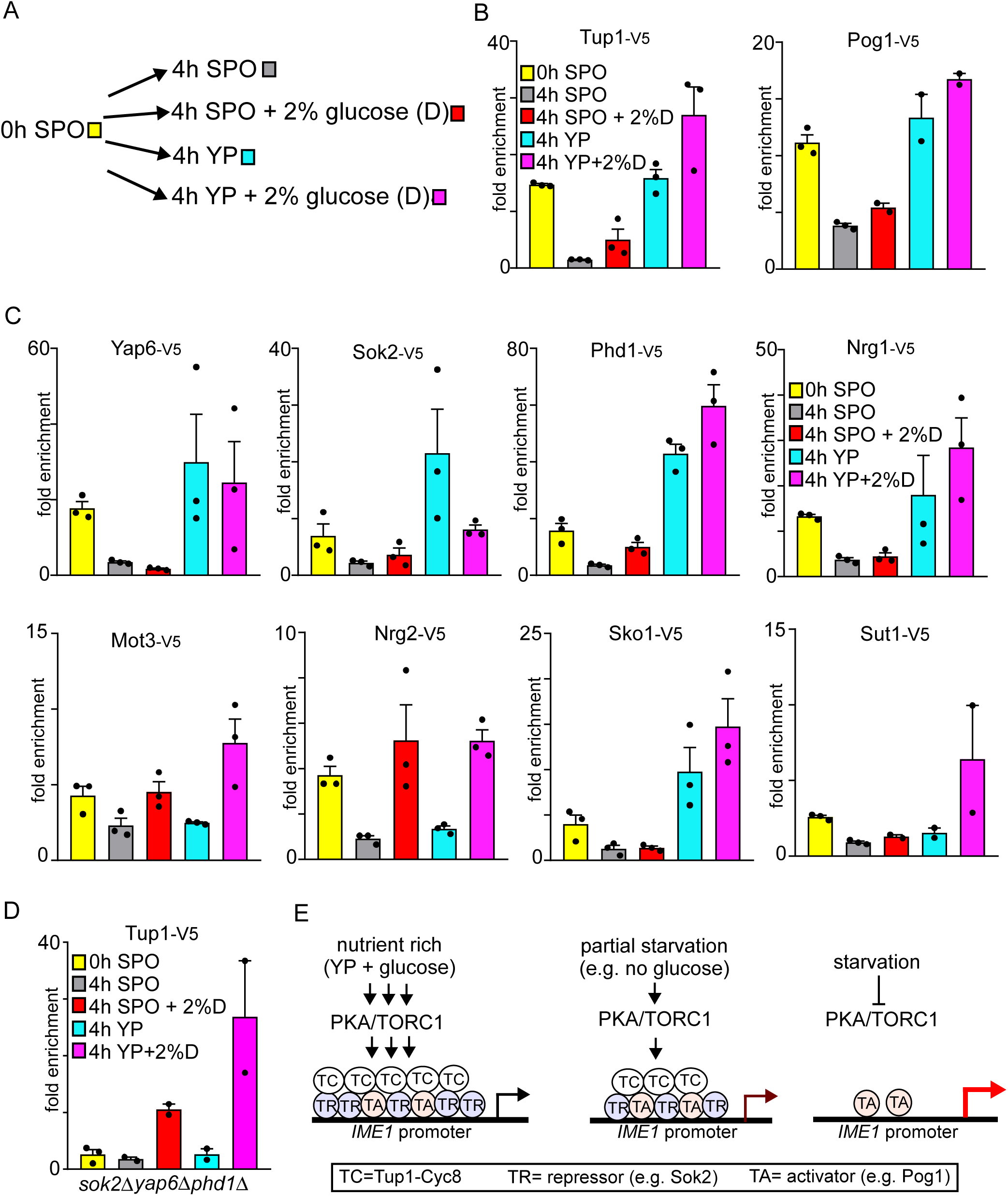
Nutrient signalling triggers distinct responses of transcription factors binding with the *IME1* promoter. (A) Scheme of experimental set up. Following growth in YPD and pre-sporulation media, cells were shifted to SPO, SPO plus 2% glucose, YP or YP plus 2% glucose for 4 hours. (B) Binding of Tup1 and Pog1 at the *IME1* promoter under distinct nutrient conditions described in A and measured by ChIP. Diploid cells harbouring V5 tagged Tup1 (FW3456) or Pog1 (FW968) were used for the analyses. Mean signals and SEM are displayed. (C) Similar to B, except that V5 tagged Yap6, Sok2, Phd1, Nrg1, Mot3, Nrg2, Sko1 and Sut1 were used for the analyses (FW3833, FW5638, FW4466, FW4393, FW4383, FW4396, FW4389, and FW6974). (D) Similar to A, except that Tup1 binding with the *IME1* promoter was determined in the *sok2*Δ*phd1*Δ*yap6*Δ cells (FW4010). (E) Model for how nutrient signalling controls the *IME1* promoter. Multiple Tup1-Cyc8 recruiting transcription factors associate in nutrient rich conditions. Upon entry into meiosis transcription factors and Tup1-Cyc8 disassociate, and consequently *IME1* transcription is induced.

Given that Sok2, Phd1, and Yap6 were strongly enriched in cells exposed to YP medium (Figure 7C), we hypothesized that Tup1-Cyc8 association with the *IME1* promoter is affected in *sok2*Δ*phd1*Δ*yap6*Δ cells in YP, but not in SPO containing glucose. We therefore examined how Tup1-Cyc8 association with *IME1* promoter was affected in *sok2*Δ*phd1*Δ*yap6*Δ cells under different nutrient conditions. Indeed, in *sok2*Δ*phd1*Δ*yap6*Δ cells, Tup1 binding was detected in SPO plus glucose, but not in YP medium (Figure 7D). These data indicate that Sok2, Phd1, and Yap6 are important for mediating Tup1-Cyc8 association in YP, while other transcription factors are required for glucose signalling to the *IME1* promoter (Figure 7E). In summary, our analyses revealed that the association of one set of transcription factors (i.e. Mot3 and Nrg2) with the *IME1* promoter is induced by glucose signalling, while another set of transcription factors (i.e. Yap6, Sok2, Phd1, Sko1, and Nrg1) was recruited to the *IME1* promoter in cells exposed to YP. Thus, only when all the required nutrient signalling pathways are repressed, all transcription factors interacting with Tup1-Cyc8 dissociate from the *IME1* promoter allowing activation of *IME1* transcription. Taken together, we propose that transcription factors important for Tup1-Cyc8 recruitment to the *IME1* promoter respond to different environmental cues to ensure tight control of *IME1* expression and thereby regulate the fate decision to enter meiosis in yeast.

## Discussion

Our study of the *IME1* promoter sheds new light on how the fate decision to enter meiosis is regulated. We report that the Tup1-Cyc8 complex together with multiple sequence-specific transcription factors constitute the essential components that control repression of the *IME1* promoter. The decision to enter meiosis and produce gametes is remarkably simple in yeast: environmental signals regulate the association and disassociation of transcription factors that recruit Tup1-Cyc8 to the *IME1* promoter. We propose that multiple transcription factors ensures repression of *IME1* expression under various environmental conditions and thus establishes tight control of entry into meiosis in yeast.

### Model for Tup1-Cyc8 mediated regulation of the *IME1* promoter

Repression of *IME1* transcription, and not activation, is highly regulated. Depletion of either Tup1 or Cyc8 completely de-repressed *IME1* expression (Figure 1). Remarkably, there was little or no delay between depletion of Tup1 and *IME1* transcription in the presence of ample nutrients. From these two observations, we can infer two important features of the *IME1* promoter. First, the transcriptional activators are bound to the *IME1* promoter or readily available prior to activation of *IME1* transcription. Second, transcriptional activators do not require nutrient or environmental signalling to activate the *IME1* promoter. Indeed, we found that the activator Pog1 is bound to the *IME1* promoter prior to activation. Pog1 does not have clear DNA binding domain, and thus Pog1 likely interacts with other proteins to associate with the *IME1* promoter. A *pog1* mutant has only a mild effect on *IME1* expression indicating that there must be other transcriptional activators controlling *IME1* transcription ^7^. Several transcriptional activators have been implicated in regulating *IME1* transcription that have not been linked with Tup1-Cyc8 ^12^.

How does Tup1-Cyc8 control the *IME1* promoter? The Tup1-Cyc8 regulates transcription of a subset of promoters in yeast ^18, 46^. Several mechanisms have been described for Tup1-Cyc8 mediated gene repression ^21, 23, 24^. For example, Tup1-Cyc8 interacts with HDACs, which in turn facilitate repression of promoters by deacetylating nucleosomes. However, we find that deletion mutants of HDACs have little effect on *IME1* expression and do not mimic *IME1* expression levels detected in Tup1 or Cyc8 depleted cells. Our data are largely consistent with a model in which Tup1-Cyc8 masks or shields activating transcription factors from recruiting co-activators at promoters ^25^. In line with Tup1-Cyc8 depletion experiments described in this work, *IME1* was among the genes that showed increased transcription upon rapid depletion of Tup1-Cyc8 from nucleus ^25^. Furthermore, we show that the transcription activator Pog1 is bound to the *IME1* promoter prior to activation, and remains bound during activation of *IME1* transcription.

Transcription factors that interact with Tup1-Cyc8 can converted into transcriptional activators in the absence of Tup1-Cyc8 ^25, 39^. We showed that multiple (at least eight) transcription factors that are known to interact with Tup1-Cyc8 associate with the *IME1* promoter. However, our data suggest these transcription factors are important for facilitating Tup1-Cyc8 binding but play little role in *IME1* transcriptional activation. Multiple lines of evidence indicate that Tup1-Cyc8 acts predominantly as a co-repressor at the *IME1* promoter. First, almost all transcription factors involved in Tup1-Cyc8 recruitment dissociate from the *IME1* promoter upon activation of *IME1* transcription (Figure 5A, SPO 4h). Second, deleting multiple transcription factors led to activation, not repression of *IME1* transcription. For example, cells lacking four of the transcription factors that showed the strongest enrichment with the *IME1* promoter (*sok2*Δ*phd1*Δ*yap6*Δ*nrg1*Δ) displayed an earlier onset of meiosis (Supplementary Figure 6). We cannot exclude that the Tup1-Cyc8 recruiting transcription factors can function as transcriptional activators in some conditions. Indeed, Yap6 and Sok2 have both been implicated as activators of transcription at some promoters ^47, 48^. In the context of the *IME1* promoter, however, transcription factors mediating Tup1-Cyc8 recruitment and transcriptional activators are likely not the same. Each transcription factor likely has a designated function in either repression or activation of *IME1* transcription. We propose that multiple transcription factors ensure that Tup1-Cyc8 co-repressor is bound to the *IME1* promoter. The Tup1-Cyc8 co-repressor complexes, in turn, mask transcriptional activators (which are different from Tup1-Cyc8 recruiting transcription factors) and prevent them from recruiting co-activators to induce *IME1* transcription.

### The regulatory logic of employing multiple sequence specific transcription factors to repress the *IME1* promoter

Why are multiple transcription factors required for recruiting Tup1-Cyc8 to the *IME1* promoter? Expression of *IME1* only occurs when the cells are starved for nitrogen and glucose ^4^. Under other environmental conditions, the *IME1* promoter must be repressed to prevent cells from inappropriately entering meiosis and form gametes. We propose that multiple transcription factors ensure *IME1* repression under various environmental conditions. First, we found that distinct sets of transcription factors associate with the *IME1* promoter in different nutrient environments (Figure 7). Second, deleting three transcription factors (Sok2, Phd1, and Yap6) led to very mild *IME1* expression in rich medium containing glucose, but *IME1* was almost fully expressed in cells grown in an acetate containing medium (Figure 4A, 4B and 5C). Thus, additional transcription factors facilitate Tup1-Cyc8 association with the *IME1* promoter in rich medium containing glucose.

How nutrient signalling controls the association of transcription factors directing Tup1-Cyc8 to the *IME1* promoter is key to dissecting how meiotic entry is regulated. We previously showed that inhibiting PKA and TORC1 is sufficient to drive entry into meiosis despite that cells were exposed to a nutrient rich medium ^16^. Here we showed that inhibiting PKA and TORC1 lowers the abundances of transcription factors important for Tup1-Cyc8 recruitment to the *IME1* promoter (Sok2, Phd1 and Yap6) (Figure 6C), however the mechanisms remains unclear. One possibility is that PKA or TORC1 phosphorylation controls the stability of Sok2, Phd1 and Yap6. Indeed, Sok2 is a direct substrate of PKA ^11, 49^. It is also possible that nutrient or stress induced phosphorylation regulates the interaction between transcription factors and Tup1-Cyc8 ^39, 50^. In addition, nutrient signalling may regulate Tup1-Cyc8 itself. Several studies have shown that sumoylation modulates the function of Tup1-Cyc8 ^51, 52^. Our data further suggest that PKA and TORC1 locally regulates the *IME1* promoter. We observed no change in cellular localization and only a minor change in abundance during meiotic entry, while chemical inhibition of PKA and TORC1 resulted in much lower levels of Sok2, Phd1 and Yap6 (Figure 6). Perhaps, PKA and TORC1 associate with *IME1* promoter directly. Future work will pinpoint the mechanism by which nutrient signalling pathways control the transcription factors regulating Tup1-Cyc8 binding with the *IME1* promoter.

### Concluding remarks

Our observation that regulated repression by multiple transcription factors controls a cell fate decision in yeast shows similarities with how multicellular organisms undertake developmental decisions. In *Drosophila*, plants, and mammals, transcriptional repressors of the Groucho family (structurally related to Tup1) are important for regulating of various developmental processes such as body patterning and determination of organ identity ^53–55^. Like Tup1-Cyc8, the association of Groucho repressor with promoters relies on sequence specific transcription factors, and Groucho repressor integrates multiple signals to control gene expression and cell fate outcomes. Regulation of the *IME1* promoter also demonstrates features of enhancer-directed transcriptional control of cell-fate master regulators in mammalian cells ^56–58^. Like the *IME1* promoter, an array of transcription factors associate and control the activity of enhancers. In addition, developmentally controlled enhancers are typically regulated by multiple upstream signalling pathways and are often primed for activation ^59^. Our findings in yeast may provide insights to better understand how signal integration controls master regulatory genes and developmental decisions in all eukaryotic cells.

## Acknowledgements

We are grateful to Andrew Wu and Fabien Moretto for their critical reading of the manuscript, and to Andreas Doncic for providing the mNeongreen tagging cassette. This work was supported by the Francis Crick Institute (FC001203), which receives its core funding from Cancer Research UK (FC001203), the UK Medical Research Council (FC001203), and the Wellcome Trust (FC001203).

## Supplementary Information

Materials and Methods

Supplementary References

Supplementary Figure legends

Supplementary Figures 1-8

Supplementary Tables 1 and 2

