## supplementary information for "Regulated repression, and not activation, governs the cell fate promoter controlling yeast meiosis"

Materials and Methods

Supplementary References

Supplementary Figure legends

Supplementary Figures 1-8

Supplementary Tables 1 and 2

#### Materials and Methods:

##### Yeast strains and plasmids

The *Saccharomyces cerevisiae* SK1 genetic background was employed for all experiments in this study. Experiments were carried out with diploid cells and the list of yeast strains described in this study can be found in Supplementary Table 1. Gene deletions, *IME1* promoter truncations, and protein fusions were achieved by the single step PCR-based gene modification protocol described in [1]. Plasmids used for gene/promoter deletions and endogenous gene tagging are summarised in Supplementary Table 2. The analogue-sensitive *tpk1-as* strains were generated by creating a M164G point mutation in the Tpk1 subunit and ablating the redundant *TPK2* and *TPK3* genes [2]. Auxin-based depletion of Tup1 and Cyc8 was achieved by fusing Tup1 and Cyc8 with the auxin-induced degron (AID) tag containing 3xV5 epitope and the *Arabidopsis thaliana* IAA7 protein [3]. The *Oryza sativa* TIR1 ligase (*osTIR1*) was also expressed under the *TEF1* promoter from a plasmid integrated at the *HIS3* locus (courtesy of Leon Chan) in the *TUP1-AID* and *CYC8-AID* strains. Protein-mNeonGreen fusions were constructed by tagging the proteins with mNeonGreen tagging cassettes (courtesy of Andreas Doncic) described in [4]. Cells expressing protein-mNeonGreen fusions also harboured a nuclear localisation signal peptide derived from simian virus 40 tagged with two copies of mCherry (2xmCherry-SV40NLS) [4].

Single-copy integration plasmids with *IME1* N-terminally fused with *sfGFP* and full length *IME1* promoter were derived from *pNH604* [5]. The *pIME1-sfGFP-IME1* fragment (~4.6 kb) was amplified from yeast cells expressing sfGFP-Ime1 and was cloned into *pNH604* plasmid into NotI and BamHI sites by restriction digestion. The resulting plasmid (*pIME1-WT*) consists of *pIME1-sfGFP-IME1* followed by *C. glabrata TRP1* and the whole cassette is flanked by the 5' and 3' UTRs of *S. cerevisiae TRP1*. To mutate transcription factor binding sites, DNA fragments of 500 bp in length corresponding to 701-1100 bp upstream of *IME1* start codon with binding site mutations were synthesised (gBlocks Gene Fragments, Integrated DNA Technologies) and cloned into the *pIME1-WT* plasmid by Gibson assembly using the NEBuilder HiFi DNA Assembly Master Mix (New England BioLabs). The *pIME1-bs*Δ plasmid carried 103 mutated nucleotides (nt) between 701-1100 bp upstream of the *IME1* start codon to disrupt Yap6, Sok2, Phd1, Mot3, Sko1, Nrg1, and Nrg2 binding sites (Supplementary Figure 4). The *pIME1-spy* plasmid was designed based on the promoter sequence in *pIME1-bs*Δ by restoring the Yap6, Sok2 and Phd1 sites (33 nt) between 701-1100 bp upstream of *IME1* start codon while other transcription factor sites remained mutated (Supplementary Figure 5). Plasmid sequences were verified by Sanger sequencing. Plasmids (*pIME1-WT, pIME1-bs*Δ and *pIME1-spy*) were linearized with PmeI, were integrated by transformation into the *TRP1* locus.

**Growth conditions**

Yeast cells were grown in YPD medium (1% yeast extract, 2% peptone, 2% glucose) supplemented with 96 µg/mL tryptophan, 24 µg/mL uracil, and 12 µg/mL adenine, grown at 30°C and liquid cultures were agitated at 300 r.p.m.. To obtain exponentially growing cells (YPD (E)) and cells grown to saturation (YPD (S)), cells were grown in YPD to saturation overnight, diluted to $OD_{600}$ = 0.2, subsequently YPD (E) cells were harvested after two to three doublings generations. YPD (S) cells were grown for 20-24 hours in YPD. To induce entry into meiosis, cell were grown overnight in YPD, shifted to pre-sporulation medium BYTA (1% yeast extract, 2% tryptone, 1% potassium acetate, 50mM potassium phthalate) at $OD_{600}$ = 0.4 for 16-18 hours, and subsequently transferred to sporulation medium SPO (0.3% potassium acetate, 0.02% raffinose, pH 7.0) at $OD_{600}$ = 1.8.

To study the responses of the transcription factors in distinct nutrient conditions in Figure 7, cells were grown in YPD and pre-sporulation medium following the standard sporulation induction protocol. Subsequently, cells were shifted to four different types of media including sporulation medium (SPO), glucose-only medium (SPO + 2% Glc), YP medium without glucose (YP + 0.05% Glc), and YPD medium (YP + 2% Glc). Yeast cells were harvested for ChIP analyses at the point of shift (0h SPO) and after four hours in the different nutrient conditions.

To study the effect of Tup1 and Cyc8 depletion on *IME1* expression (Figure 1, Figure 3, Figure S2 and Figure S4), 500µM of indole-3-acetic acid (3-IAA) (Aldrich) was added to exponentially growing cells to induce degradation of Tup1-AID and Cyc8-AID proteins. As control, same volume of dimethyl sulphoxide (DMSO) was added to yeast cells. Cells were harvested at the indicated time points for ChIP, RT-qPCR, smFISH, and western blot analyses.

To inhibit PKA and TORC1 in *tpk1-as* strains in Figure 6, yeast cells were grown to saturation in YPD and treated with 1NM-PP1 (Calbiochem, Merck Millipore) and rapamycin (Sigma). Untreated cells were diluted to $OD_{600}$ = 0.1. To inhibit PKA, 5µM of 1NM-PP1 was added to cells diluted to $OD_{600}$ = 1. To inhibit TORC1, 1µg/mL rapamycin was added to cells diluted to $OD_{600}$ = 0.5. Cells treated with both inhibitors were diluted to $OD_{600}$ = 2. Yeast cultures were treated for 6 hours at 30°C, 300 r.p.m. and fixed with 5% trichloroacetic acid for Western blotting.

**Chromatin immunoprecipitation (ChIP)**

Harvested cells were crosslinked with 1% formaldehyde for 20 minutes at room temperature and reaction was quenched by the addition of 100mM glycine. Cells were washed with FA lysis buffer (50mM HEPES pH 7.5, 150mM sodium chloride, 1mM EDTA pH 7.6, 1% Triton X-100, 0.1% sodium deoxycholate, 0.1% SDS), snap frozen and stored at -80°C. Cell lysis was performed in cold FA lysis buffer with Complete Mini Protease Inhibitor Cocktail (Roche). Samples were homogenised with zirconia beads (BioSpec) using Mini-Beadbeater-96 (BioSpec). The chromatin fraction was subjected to shearing by sonication on Bioruptor Plus (Diagenode) using 9 cycles of 30s on, 30s off. V5 epitope-tagged proteins were immunoprecipitated with anti-V5 agarose beads (Sigma-Aldrich) at room temperature for two hours with rotation. Subsequently, the agarose beads were washed with FA lysis buffer, FA lysis buffer with 260mM sodium chloride, and a lithium chloride/detergent buffer (10mM Tris pH 8, 250mM lithium chloride, 0.5% NP-40, 0.5% sodium deoxycholate, 1mM EDTA). Samples were reverse crosslinked in TE buffer with 1% SDS at 65°C, 500 r.p.m. overnight and treated with 80µg/mL proteinase K (Thermo Scientific) at 37°C for two hours. Purified DNA fragments were quantified by quantitative PCR using EXPRESS SYBR GreenER SuperMix (Thermo Fisher Scientific) on Applied Biosystems 7500 Fast Real-Time PCR System (Thermo Fisher Scientific). ChIP signals were normalised over the silent mating type cassette *HMR*. Primer sequences are listed in Supplementary Table 2.

#### RNA isolation and reverse transcription

Total RNA was extracted from harvested cells using the hot phenol method. Briefly, TES buffer (10mM Tris-HCl pH 7.5, 10mM EDTA, 0.5% SDS) and acid-phenol:chloroform (Ambion) were added to samples and incubated at 65°C. RNA was subsequently purified using the NucleoSpin RNA kit (Macherey-Nagel) according to the manufacturer's instructions. rDNase was added to remove residual genomic DNA during the purification procedure. For reverse transcription, ProtoScript II First Strand cDNA Synthesis Kit (New England BioLabs) was used and 500ng of total RNA was provided as template in each reaction. qPCR reactions were prepared using EXPRESS SYBR GreenER SuperMix (Thermo Fisher Scientific) or PowerUp SYBR Green Master Mix (Thermo Fisher Scientific) and *IME1* level was quantified on Applied Biosystems 7500 Fast Real-Time PCR System (Thermo Fisher Scientific). Signals were normalised over *ACT1*. Primer sequences are listed in Supplementary table 2.

#### Western blotting

Proteins were extracted from cells fixed with 5% trichloroacetic acid by lysing cells in protein breakage buffer (50mM Tris at pH 7.5, 1mM EDTA, 27.5mM DTT) with 0.5mm glass beads (BioSpec) on the Mini-Beadbeater-96 (BioSpec). Samples were denatured in SDS loading buffer (62.5mM Tris (pH 6.8), 2% β-mercaptoethanol, 10% glycerol, 3% SDS, and 0.017% Bromophenol Blue) and separated by SDS-PAGE in Tris-glycine buffer (25mM Tris base, 192mM glycine, 0.1% SDS). V5-epitope tagged proteins were detected using anti-V5 primary antibodies (Invitrogen, 1:2000, mouse) and IRDye 800CW (LI-COR, 1:15000) or HRP-conjugated (GE Healthcare, 1:8000) secondary antibodies. For equal loading Hxk1 protein levels were determined using anti-hexokinase primary antibodies (Stratech Scientific, 1:2000, rabit), and IRDye 680RD (LI-COR, 1:15000) or HRP-conjugated (GE Healthcare, 1:8000) secondary antibodies. Images were acquired on the Odyssey CLx imaging system (LI-COR) or by ECL Prime (GE Healthcare) using Amersham Imager 600 (GE Healthcare). V5-tagged protein levels were normalised over Hxk1 signals and quantification were carried out in the Image Studio Lite software (LI-COR).

**Nuclei/DAPI counting**

Samples were fixed in 80% ethanol and stained with 1µg/mL 4',6-diamidino-2-phenylindole (DAPI) in PBS buffer. Cells with two, three or four DAPI masses were considered meiosis, while cells harbouring one DAPI mass were counted as NO meiosis. At least 200 cells were analysed for each sample.

**Microscopy**

For single molecule RNA fluorescence *in situ* hybridisation (smFISH), cells were fixed with formaldehyde and washed with Buffer B (1.2M sorbitol, 0.1M potassium phosphate dibasic, pH 7.5). To spheroplast the cells, samples were treated with 40µg/mL Zymolyase-100T (MP Biomedicals) and 57.2mM β-mercaptoethanol in Buffer B at 30°C. Spheroplasted cells were washed and smFISH probes recognising *IME1* (AF594) and *ACT1* (Cy5) [2] were hybridised overnight at 30C. Cells were washed with wash buffer, stained with 1µg/mL DAPI in wash buffer and re-suspended 2XSSC. Subsequently, cells were spun down and resuspended prior imaging in anti-fade GLOX with 1% catalase (Sigma) and 1% glucose oxidase (Sigma) [6] prior imaging. Images were acquired using the Eclipse Ti-E inverted microscope system (Nikon) using the 100x oil objective with the ORCA-FLASH 4.0

camera (Hamamatsu) and NIS-elements software (Nikon). Cells were imaged at every 0.3µm along the z-axis using the built-in z-axis drive and a total of 25 images were taken for each z-stack. Signals from all the planes were merged into a 2D image by applying maximum intensity z-projection in ImageJ [7]. Only cells with *ACT1* signals were considered for *IME1* quantification using the StarSearch software (ttp://rajlab.seas.upenn.edu/StarSearch/launch.html).

For the quantification of sfGFP expressed in *pIME1-WT* and *pIME1-bs*Δ presented in Figure 3D, cells were induced to sporulate using the standard protocol and samples were taken from the sporulation culture at indicated time points. Harvested cells were fixed with formaldehyde and re-suspended prior in a buffer containing 16.6mM potassium phosphate monobasic, 83.4mM potassium phosphate dibasic, and 1.2M sorbitol for imaging. Imaging was carried out on the Eclipse Ti-E inverted microscope system (Nikon) using the 100x oil objective with the ORCA-FLASH 4.0 camera (Hamamatsu) and NIS-elements software (Nikon). sfGFP signals in each cell were quantified using the ImageJ software [7].

For determining the localization of transcription factors tagged with mNeongreen described in Figure 6B and Supplementary Figure 8, cells were induced to sporulate with the standard protocol. Subsequently, cells were imaged at 0h in SPO and 4h in SPO. Images were acquired using the same imaging system and set up described for quantification of sfGFP expressed in *pIME1-WT* and *pIME1-bs*Δ. Signals from whole cell and cell nucleus were quantified with ImageJ software and with use of the nuclear marker (2xmCherry-SV40NLS) [7]. Signal from cytosol was inferred from the difference between whole cell signal and nucleus signal.

#### *IME1* promoter motif analysis

Predicted transcription factor binding sites by scanning the *IME1* promoter with the curated transcription factor motifs in the YeTFaSCo database [8]. Binding sites were predicted to have at least 75% of the maximum possible score, with the exception of Sut1 binding site which was predicted with a 70% threshold. The Sko1 binding site was identified manually by scanning the promoter sequencing and compare to the consensus motif TGACG as described in [8].

#### Statistical analyses

Data statistics and statistical analyses indicated in the figure legends were computed using GraphPad Prism version 8.2.0 for Windows, GraphPad Software, San Diego, California USA, www.graphpad.com. Data from the smFISH experiments presented in Figure 1H, 4A, and 4D were analysed using unpaired parametric two-tailed Welch's t-test with 95% confidence. P-values are indicated in the figures, where ns stands for non-significant, * = ≤ 0.05, ** = ≤ 0.01, *** = ≤ 0.001.

#### Supplementary Figure legends:

**Supplementary Figure 1.**

The effect of truncations in the *IME1* promoter on meiosis. For the analyses we used diploid cells harbouring one copy of *IME1* deleted (control, FW4128), while different promoter deletion mutants were generated at the wild-type *IME1* locus (*pIME1*(-1350-2315∆), FW4781; *pIME1*(-1250-2315∆), FW4780; *pIME1*(-950-2315∆), FW4779; *pIME1*(-900-2315∆), FW4778; *pIME1*(-850-2315∆), FW4777; *pIME1*(-800-2315∆), FW3944). Control and mutant cells were grown to saturation in rich medium (YPD), grown for an additional 16 hours in pre-sporulation medium, and subsequently cells shifted to sporulation medium. Samples were taken at the indicated time points, fixed, and DAPI masses were counted for the indicated time points to determine the percentage of cells that underwent meiosis (MI+MII). Cell harbouring two, three or four DAPI masses were classified as meiosis. At least 200 cells were analysed per sample.

**Supplementary Figure 2.**

(A) Representative images of smFISH data described in Figure 1G. Cells were fixed, and hybridized with *IME1* (AF594) and *ACT1* (Cy5) probes. Representative images representing bright field, *IME1*, *ACT1*, and DAPI are displayed. (B) Same data as described in Figure 1H, except that the single cell data for *IME1* expression in *TUP1-AID*+DMSO were binned according to expression levels. (C) Same data as in Figure 1J, except that the single cell data for *IME1* were binned according to expression levels.

**Supplementary Figure 3.**

(A) Sequence of the *IME1* promoter spanning a region between 600 and 1200 bp upstream of *IME1* AUG. Highlighted are the transcription factors binding motifs in the *IME1* promoter. (B) Table describing the sequence motifs identified in the *IME1* promoter. Data are shown for transcription factors that associate with the *IME1* promoter as shown in Figure 2B, and for the region between 700 and 1100 bp upstream of *IME1* AUG. The number of binding sites as well as whether the transcription factor is known to interact with Tup-Cyc8 are displayed.

**Supplementary Figure 4.**

(A) Western blot analysis of samples described in Figure 4C. Diploid cells harbouring *TUP1-AID* and V5-tagged transcription factors (*YAP6-V5*, FW4214; *SOK2-V5*, FW4218; *PHD1-V5*, FW5056; *MOT3-V5*, FW4229; *NRG1-V5*, FW4230; *NRG2-V5*, FW5055; *SKO1-V5*, FW4224)were grown to exponential phase. As a control *TUP1-AID* cells (FW5057) were included. The *TUP1-AID* also harbours a V5 tag. Cells were either treated with IAA (+) or DMSO (-), and the expression for each transcription factor and *TUP1* was determined by western blot using anti-V5 antibodies. Highlighted are the bands representing the different transcription factors. (B) Sequence displaying the mutated sites (in red lower case) in the *IME1* promoter of *pIME1-bs*Δ. (C) Meiosis in cells harbouring *pIME1-WT* (FW5370) and *pIME1-bs*Δ ( FW5372). Samples were taken at the indicated time points, fixed, and DAPI masses were counted for the indicated time points to determine the percentage of cells that underwent meiosis (MI+MII). Cell harbouring two, three or four DAPI masses were classified as meiosis.

**Supplementary Figure 5.**

Sequence displaying the wild-type motif sites (in blue uppercase) in the *IME1* promoter of *pIME1-spy.*

**Supplementary Figure 6.**

Meiosis in wild-type (FW3456), *sok2∆phd1∆yap6∆* triple mutant (FW4010), and *sok2∆phd1∆yap6∆nrg1∆* quadruple mutant cells (FW5657). Cells were induced to enter meiosis in SPO. DAPI masses were counted for the indicated time points to determine the percentage of cells that underwent meiosis (MI+MII). Cell harbouring two, three or four DAPI masses were classified as meiosis. At least 200 cells were analysed.

**Supplementary Figure 7.**

Binding of Sok2, Phd1, Yap6, Tup1 and Cyc8 under different nutrient conditions to the *IME1* promoter as determined by ChIP. Cells harbouring V5 epitope tagged version of each transcription factor (Yap6-V5, FW3833; Sok2-V5, FW5638; Phd1-V5, FW4466; Tup1-V5, FW3456; Cyc8-V5, FW6381) were grown till exponential growth (E) or saturation (S) in YPD, or grown for an additional 18 hours in pre-sporulation (0h) or shifted to sporulation medium (4h). ChIP signals were normalized over *HMR*. The mean and SEM are displayed. The data complement Figure 6A.

**Supplementary Figure 8.**

(A) Representative images of Sok2-mNG (FW7475), Phd1-mNG (FW7477), Yap6-mNG (FW7473), Tup1-mNG (FW7644), and Cyc8-mNG (FW7642) localization prior (0 hours in SPO), and during entry into meiosis (4 hours in SPO). Each transcription factor was fused to mNeongreen (mNG). These cells also expressed a mCherry fused to SV40 nuclear localization signal (NLS) (mCherry-NLS) to determine the nuclear localization. (B) Quantification of nuclear signals of transcriptional factors described in A. As a control, the signals of cells harbouring no mNG-tag (FW5199) are displayed. The black bar indicates the mean signal, and each point displays a single cell measurement. The error bars represent the 95% confidence interval.

### Supplementary Figure 1

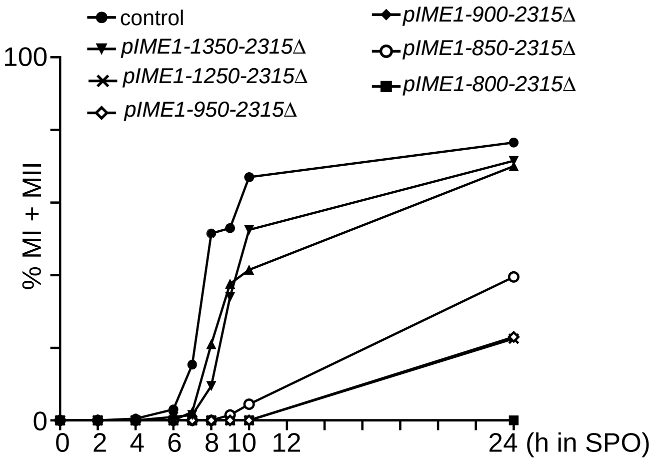

Supplementary Figure 2

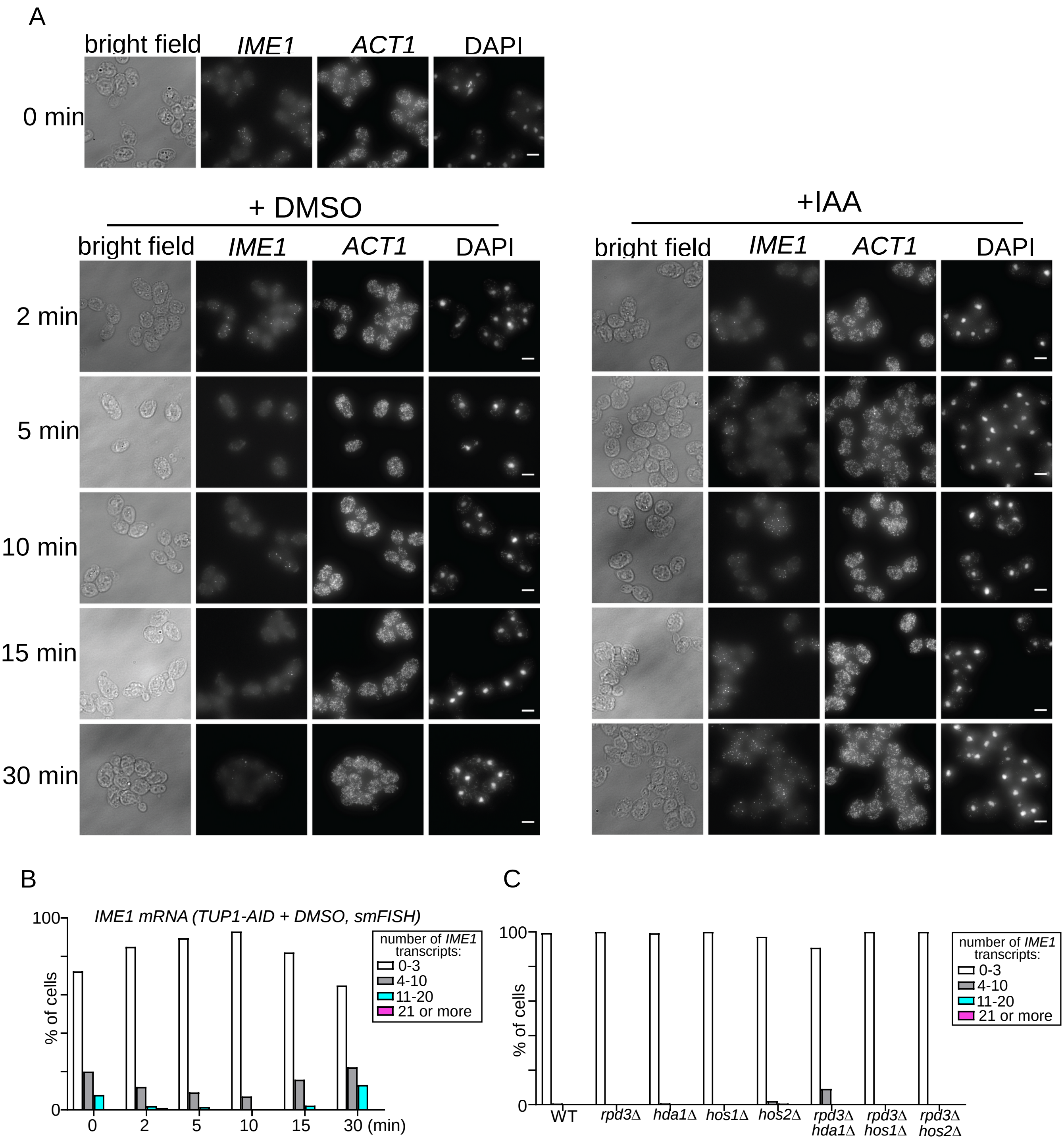

Supplementary Figure 3

A

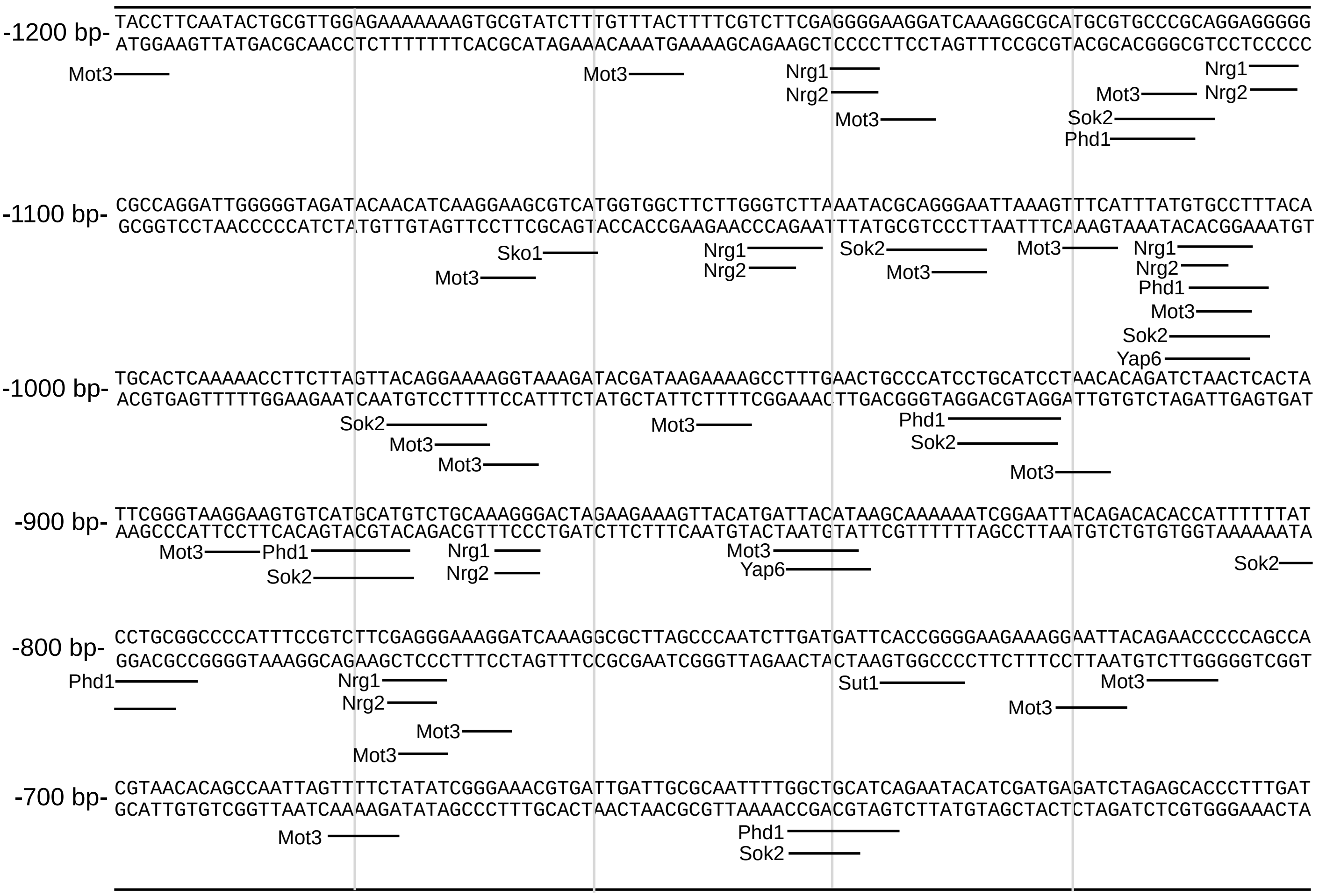


B

| Putative BS in -700 / -1100 | | Interacts with Tup1-Cyc8 |
|---|---|---|
| Yap6 | 2x | y |
| Sok2 | 6x | nd |
| Phd1 | 4x | y |
| Nrg1 | 4x | y |
| Nrg2 | 4x | y |
| Sut1 | 1x | y |
| Mot3 | 14x | y |
| Sko1 | 1x | y |

Supplementary Figure 4

A

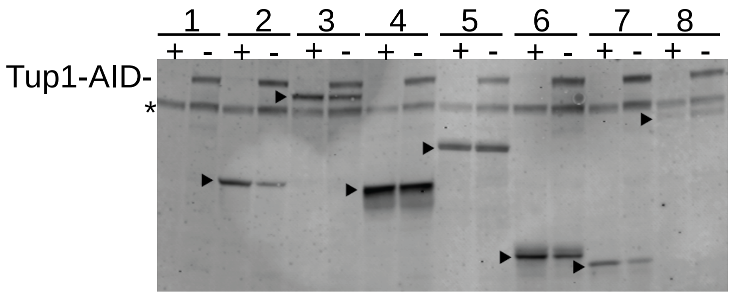


1. Tup1-AID
2. Tup1-AID + Yap6-V5
3. Tup1-AID + Sok2-V5
4. Tup1-AID + Phd1-V5
5. Tup1-AID + Mot3-V5
6. Tup1-AID + Nrg1-V5
7. Tup1-AID + Nrg2-V5
8. Tup1-AID + Sko1-V5

B

*pIME-bsΔ*

-1100 bp- CGCCAGGATTGGGGGTAGATACAACATCAAttcAGatgacTGGTGGCTTCTTttttTCTTAAATACtactttcATTAAAGggTgATTTAgtgtaagTTcac
GCGGTCCTAACCCCCATCTATGTTGTAGTTaagTCtactgACCACCGAAGAAaaaaAGAATTTATGatgaaagTAATTTCccAcTAAATcacattcAAgtg

-1000 bp- gtacCTCAAAAACCTTCTTAGTTACcttccActtTcAAGATACGATAAtAccAGCCTTTGAACTGCCCATCagtacgaaTAACACAGATCTAACTCACTA
catgGAGTTTTTGGAAGAATCAATGgaaggTgaaAgTTCTATGCTATTaTggTCGGAAACTTGACGGGTAGtcatgcttATTGTGTCTAGATTGAGTGAT

-900 bp- TTCGGGTAAttcAGTGTCAgtacTGTagtaAActttACTAGAAGAAAGTTACATGAggcacgccGCAAAAAATCGGAATTACAGACACACCATTTTTTAT
AAGCCCATTaagTCACAGTcatgACAtcatTTgaaaTGATCTTCTTTCAATGTACTccgttgtcCGTTTTTTAGCCTTAATGTCTGTGTGGTAAAAAATA

-800 bp- aagtaGtaCCCATTTCCGTCTTCGctttcAAttcTCAAAGGCGCTTAGCCCAATCTTGATGATTCACCGGGGAAGAAAttcATgcaAtAcaaCCCAGCCA
ttcatCatGGGTAAAGGCAGAAGCcaaagTTaagAGTTTCCGCGAATCGGGTTAGAACTACTAAGTGGCCCCTTCTTTaagTAcgtTaTgttGGGTCGGT

*pIME-spy*

```
-1100 bp- CGCCAGGATTGGGGGTAGATACAACATCAAttcAGatgacTGGTGGCTTCTTtttTCTTAAATACGCAGttcATTAAAGggTgATTTAgtgtaagTTcac
          GCGGTCCTAACCCCCATCTATGTTGTAGTTaagTCtactgACCACCGAAGAAaaaAGAATTTATGCGTCaagTAATTTCccAcTAAATcacattcAAgtg

-1000 bp- TGCACTCAAAAACCTTCTTAGTTACcttccActtTcAAGATACGATAAtAccAGCCTTTGAACTGCCCATCCTGCAgaaTAACACAGATCTAACTCACTA
          ACGTGAGTTTTTGGAAGAATCAATGgaaggTgaaAgTTCTATGCTATTaTggTCGGAAACTTGACGGGTAGGACGTcttATTGTGTCTAGATTGAGTGAT

-900 bp- TTCGGGTAAttcAGTGTCAgtacTGTCTGCAAccttACTAGAAGAAAGTTACATGATTACATAAGCAAAAAATCGGAATTACAGACACACCATTTTTTAT
         AAGCCCATTaagTCACAGTcatgACACACGTTgaaaTGATCTTCTTTCAATGTACTAATGTATTCGTTTTTTAGCCTTAATGTCTGTGTGGTAAAAAATA

-800 bp- aCTGCGtaCCCATTTCCGTCTTCGctttcAAttcTCAAAGGCGCTTAGCCCAATCTTGATGATTCACCGGGGAAGAAAttcATgcaAtAcaaCCCAGCCA
         tGACGCatGGGTAAAGGCAGAAGCcaaagTTaagAGTTTCCGCGAATCGGGTTAGAACTACTAAGTGGCCCCTTCTTTaagTAcgtTaTgttGGGTCGGT
```

Supplementary Figure 6

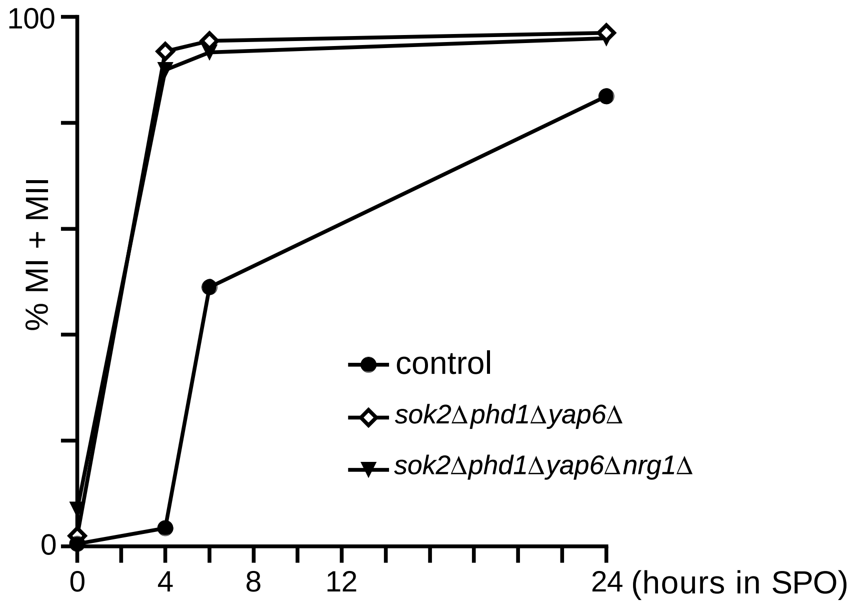

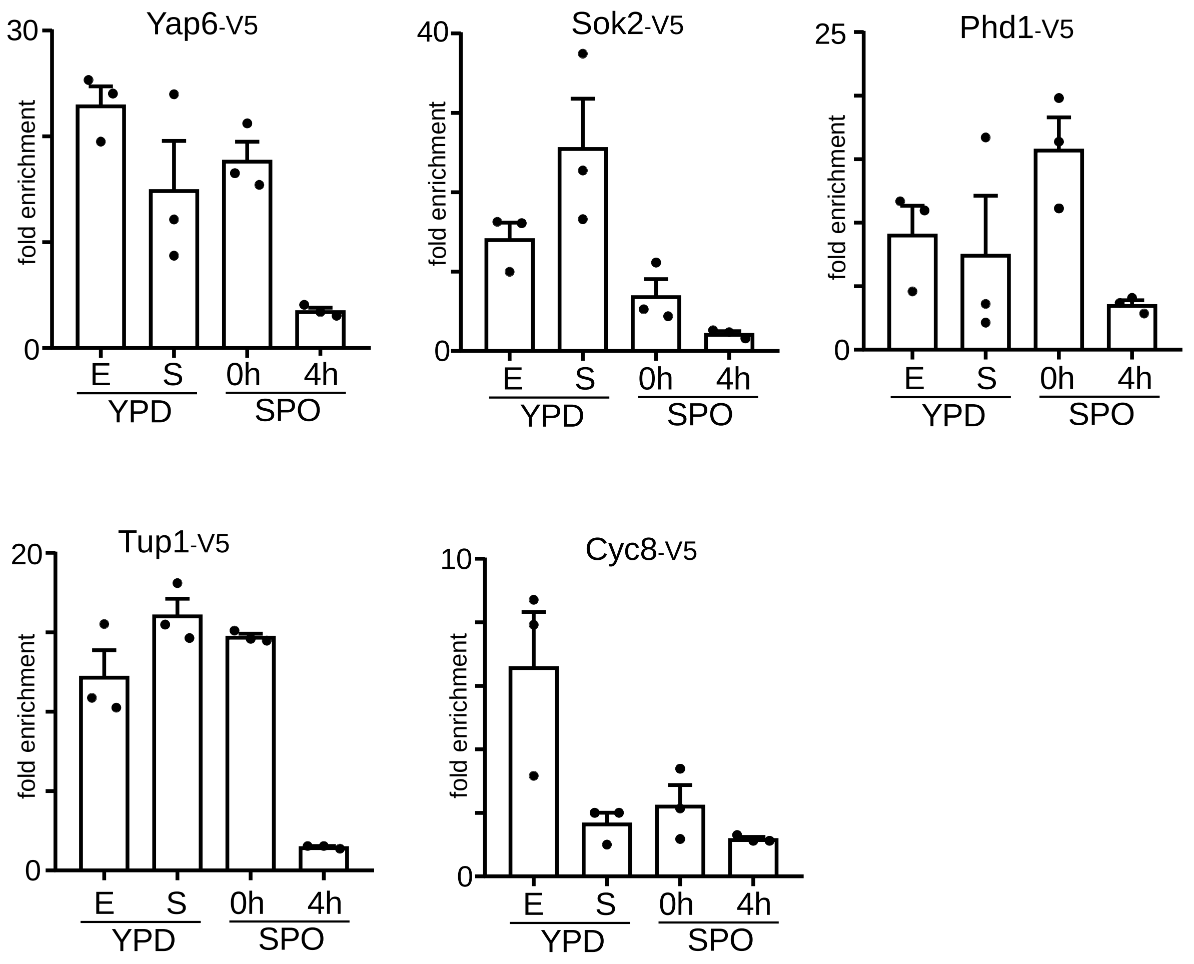
Yap6-V5
30
0
fold enrichment
E
S
0h
4h
YPD
SPO
Sok2-V5
40
0
fold enrichment
E
S
0h
4h
YPD
SPO
Phd1-V5
25
0
fold enrichment
E
S
0h
4h
YPD
SPO
Tup1-V5
20
0
fold enrichment
E
S
0h
4h
YPD
SPO
Cyc8-V5
10
0
fold enrichment
E
S
0h
4h
YPD
SPO

Supplementary Figure 8

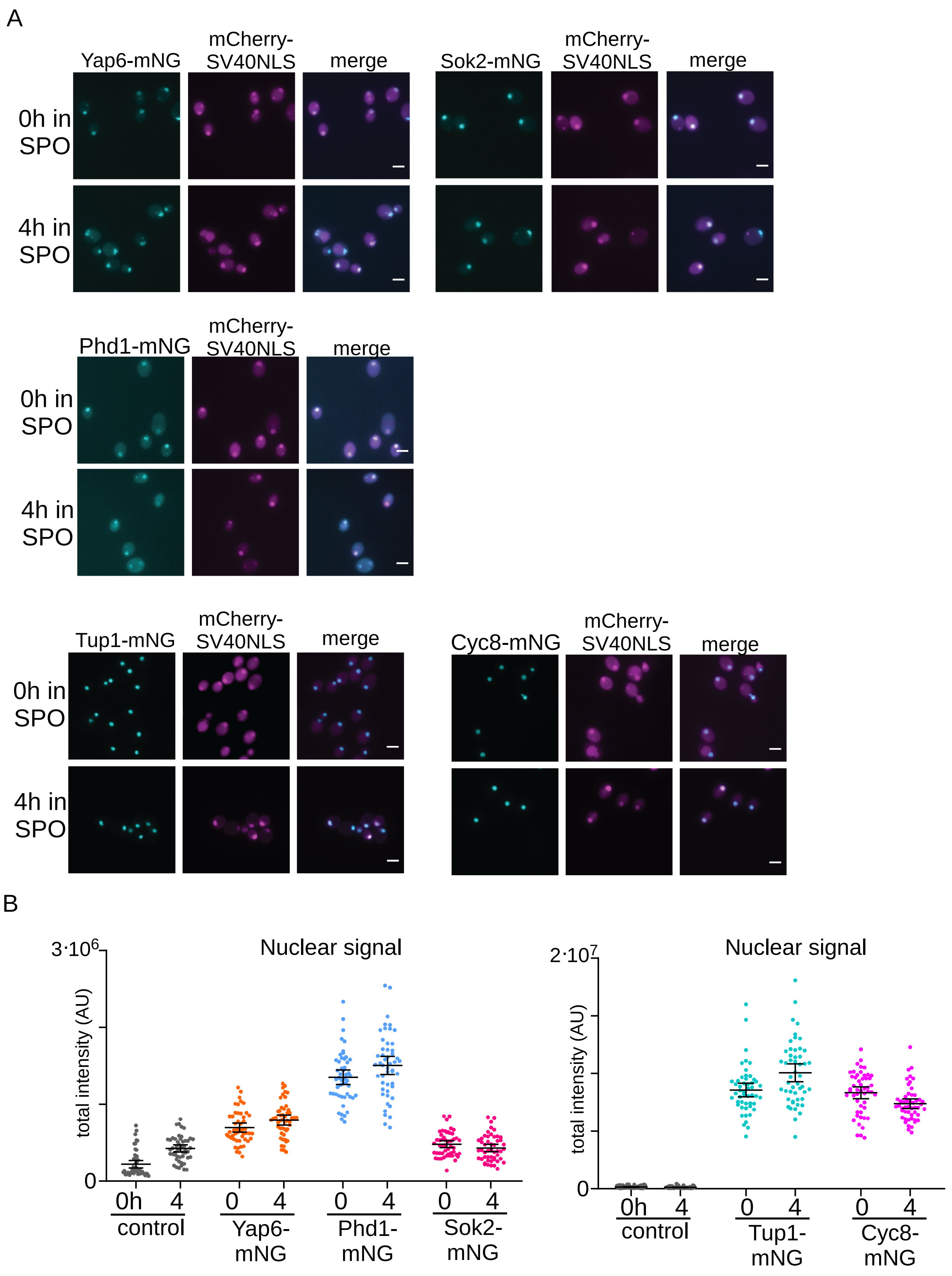

**Supplementary Table 1. Genotypes of strains used througout this study**

| Yeast strain | Genotype |
|---|---|
| FW1511 (*WT*) | MATa, ho::LYS2, lys2, ura3, leu2::hisG, his3::hisG, trp1::hisG<br>MATalpha, ho::LYS2, lys2, ura3, leu2::hisG, his3::hisG, trp1::hisG |
| FW4128 | MATa, ho::LYS2, lys2, ura3, leu2::hisG, his3::hisG, trp1::hisG<br>MATalpha, ho::LYS2, lys2, ura3, leu2::hisG, his3::hisG, trp1::hisG, ime1::HISMX6 |
| FW3946 | MATa, ho::LYS2, lys2, ura3, leu2::hisG, his3::hisG, trp1::hisG, pIME1(-1600-2315)::KanMX<br>MATalpha, ho::LYS2, lys2, ura3, leu2::hisG, his3::hisG, trp1::hisG, ime1::HISMX6 |
| FW3947 | MATa, ho::LYS2, lys2, ura3, leu2::hisG, his3::hisG, trp1::hisG, pIME1(-1400-2315)::KanMX<br>MATalpha, ho::LYS2, lys2, ura3, leu2::hisG, his3::hisG, trp1::hisG, ime1::HISMX6 |
| FW3948 | MATa, ho::LYS2, lys2, ura3, leu2::hisG, his3::hisG, trp1::hisG, pIME1(-1200-2315)::KanMX<br>MATalpha, ho::LYS2, lys2, ura3, leu2::hisG, his3::hisG, trp1::hisG, ime1::HISMX6 |
| FW3949 | MATa, ho::LYS2, lys2, ura3, leu2::hisG, his3::hisG, trp1::hisG, pIME1(-1000-2315)::KanMX<br>MATalpha, ho::LYS2, lys2, ura3, leu2::hisG, his3::hisG, trp1::hisG, ime1::HISMX6 |
| FW3950 | MATa, ho::LYS2, lys2, ura3, leu2::hisG, his3::hisG, trp1::hisG, pIME1(-800-2315)::KanMX<br>MATalpha, ho::LYS2, lys2, ura3, leu2::hisG, his3::hisG, trp1::hisG, ime1::HISMX6 |
| FW3951 | MATa, ho::LYS2, lys2, ura3, leu2::hisG, his3::hisG, trp1::hisG, pIME1(-600-2315)::KanMX<br>MATalpha, ho::LYS2, lys2, ura3, leu2::hisG, his3::hisG, trp1::hisG, ime1::HISMX6 |
| FW6381 | MATa, ho::LYS2, lys2, ura3, leu2::hisG, his3::hisG, trp1::hisG, CYC8-3V5::KanMX<br>MATalpha, ho::LYS2, lys2, ura3, leu2::hisG, his3::hisG, trp1::hisG, CYC8-3V5::KanMX |
| FW3944 | MATa, ho::LYS2, lys2, ura3, leu2::hisG, his3::hisG, trp1::hisG, pIME1(-800-2315)::HIS3MX<br>MATalpha, ho::LYS2, lys2, ura3, leu2::hisG, his3::hisG, trp1::hisG, ime1::HISMX6 |
| FW4777 | MATa, ho::LYS2, lys2, ura3, leu2::hisG, his3::hisG, trp1::hisG, pIME1(-850-2315)::HIS3MX<br>MATalpha, ho::LYS2, lys2, ura3, leu2::hisG, his3::hisG, trp1::hisG, ime1::HISMX6 |
| FW4778 | MATa, ho::LYS2, lys2, ura3, leu2::hisG, his3::hisG, trp1::hisG, pIME1(-900-2315)::HIS3MX<br>MATalpha, ho::LYS2, lys2, ura3, leu2::hisG, his3::hisG, trp1::hisG, ime1::HISMX6 |
| FW4779 | MATa, ho::LYS2, lys2, ura3, leu2::hisG, his3::hisG, trp1::hisG, pIME1(-950-2315)::HIS3MX<br>MATalpha, ho::LYS2, lys2, ura3, leu2::hisG, his3::hisG, trp1::hisG, ime1::HISMX6 |
| FW4780 | MATa, ho::LYS2, lys2, ura3, leu2::hisG, his3::hisG, trp1::hisG, pIME1(-1250-2315)::HIS3MX<br>MATalpha, ho::LYS2, lys2, ura3, leu2::hisG, his3::hisG, trp1::hisG, ime1::HISMX6 |
| FW4781 | MATa, ho::LYS2, lys2, ura3, leu2::hisG, his3::hisG, trp1::hisG, pIME1(-1350-2315)::HIS3MX<br>MATalpha, ho::LYS2, lys2, ura3, leu2::hisG, his3::hisG, trp1::hisG, ime1::HISMX6 |
| FW5057 | MATa, ho::LYS2, lys2, ura3, leu2::hisG, trp1::hisG, his3::pTEF1-osTIR::HIS3, TUP1-3V5-IAA7::KanMX6<br>MATalpha, ho::LYS2, ura3, leu2::hisG, his3::hisG, trp1::hisG, his3::pTEF1-osTIR::HIS3, TUP1-3V5-IAA7::KanMX6 |
| FW6371 | MATa, ho::LYS2, lys2, ura3, leu2::hisG, his3::hisG, trp1::hisG, his3::pTEF1-osTIR::HIS3 , CYC8-3V5-IAA7::KanMX6<br>MATalpha, ho::LYS2, lys2, ura3, leu2::hisG, his3::hisG, trp1::hisG, his3::pTEF1-osTIR::HIS3, CYC8-3V5-IAA7::KanMX6 |
| FW3456 | MATa, ho::LYS2, lys2, ura3, leu2::hisG, his3::hisG, trp1, TUP1-3V5::HIS3<br>MATalpha, ho::LYS2, lys2, ura3, leu2::hisG, his3::hisG, trp1, TUP1-3V5::HIS3 |
| FW968 | MATa, ho::LYS2, ura3, leu2::hisG, his3::hisG, trp1::hisG , POG1-3xV5::HIS3<br>MATalpha, ho::LYS2, ura3, leu2::hisG, his3::hisG, trp1::hisG , POG1-3xV5::HIS3 |
| FW3833 | MATa, ho::LYS2, lys2, ura3, leu2::hisG, his3::hisG, trp1::hisG, YAP6-3V5::KanMX<br>MATalpha, ho::LYS2, lys2, ura3, leu2::hisG, his3::hisG, trp1::hisG, YAP6-3V5::KanMX |
| FW4466 | MATa, ho::LYS2, lys2, ura3, leu2::hisG, his3::hisG, trp1::hisG, PHD1-3V5::KanMX<br>MATalpha, ho::LYS2, lys2, ura3, leu2::hisG, his3::hisG, trp1::hisG, PHD1-3V5::KanMX |
| FW4393 | MATa, ho::LYS2, lys2, ura3, leu2::hisG, his3::hisG, trp1::hisG, NRG1-3V5::KanMX<br>MATalpha, ho::LYS2, lys2, ura3, leu2::hisG, his3::hisG, trp1::hisG, NRG1-3V5::KanMX |
| FW4396 | MATa, ho::LYS2, lys2, ura3, leu2::hisG, his3::hisG, trp1::hisG, NRG2-3V5::KanMX<br>MATalpha, ho::LYS2, lys2, ura3, leu2::hisG, his3::hisG, trp1::hisG, NRG2-3V5::KanMX |
| FW7072 | MATa, ho::LYS2, lys2, ura3, leu2::hisG, his3::hisG, trp1::hisG, CIN5-3V5::KanMX<br>MATalpha, ho::LYS2, lys2, ura3, leu2::hisG, his3::hisG, trp1::hisG, CIN5-3V5::KanMX |
| FW4665 | MATa, ho::LYS2, lys2, ura3, leu2::hisG, his3::hisG, trp1::hisG, MIG1-3V5::KanMX<br>MATalpha, ho::LYS2, lys2, ura3, leu2::hisG, his3::hisG, trp1::hisG, MIG1-3V5::KanMX |
| FW6974 | MATa, ho::LYS2, lys2, ura3, leu2::hisG, his3::hisG, trp1::hisG, SUT1-3V5::KanMX<br>MATalpha, ho::LYS2, lys2, ura3, leu2::hisG, his3::hisG, trp1::hisG, SUT1-3V5::KanMX |
| FW4383 | MATa, ho::LYS2, lys2, ura3, leu2::hisG, his3::hisG, trp1::hisG, MOT3-3V5::KanMX<br>MATalpha, ho::LYS2, lys2, ura3, leu2::hisG, his3::hisG, trp1::hisG, MOT3-3V5::KanMX |
| FW4389 | MATa, ho::LYS2, lys2, ura3, leu2::hisG, his3::hisG, trp1::hisG, SKO1-3V5::KanMX<br>MATalpha, ho::LYS2, lys2, ura3, leu2::hisG, his3::hisG, trp1::hisG, SKO1-3V5::KanMX |
| FW7070 | MATa, ho::LYS2, lys2, ura3, leu2::hisG, his3::hisG, trp1::hisG, SFL1-3V5::KanMX<br>MATalpha, ho::LYS2, lys2, ura3, leu2::hisG, his3::hisG, trp1::hisG, SFL1-3V5::KanMX |
| FW4386 | MATa, ho::LYS2, lys2, ura3, leu2::hisG, his3::hisG, trp1::hisG, RGT1-3V5::KanMX<br>MATalpha, ho::LYS2, lys2, ura3, leu2::hisG, his3::hisG, trp1::hisG, RGT1-3V5::KanMX |
| FW5638 | MATa, ho::LYS2, ura3, leu2::hisG, his3::hisG, trp1::hisG, SOK2-3V5::HIS3<br>MATalpha, ho::LYS2, ura3, leu2::hisG, his3::hisG, trp1::hisG, SOK2-3V5::HIS3 |
| FW4399 | MATa, ho::LYS2, lys2, ura3, leu2::hisG, his3::hisG, trp1::hisG, SKN7-3V5::KanMX<br>MATalpha, ho::LYS2, lys2, ura3, leu2::hisG, his3::hisG, trp1::hisG, SKN7-3V5::KanMX |
| FW8102 | MATa, ho::LYS2, lys2, ura3, leu2::hisG, his3::hisG, trp1::hisG, rpd3::HIS3MX, TUP1-3V5::HIS3<br>MATalpha, ho::LYS2, lys2, ura3, leu2::hisG, his3::hisG, trp1::hisG, rpd3::HIS3MX, TUP1-3V5::HIS3 |
| FW8426 | MATa, ho::LYS2, lys2, ura3, leu2::hisG, his3::hisG, trp1::hisG, hda1::KanMX, TUP1-3V5::HIS3<br>MATalpha, ho::LYS2, lys2, ura3, leu2::hisG, his3::hisG, trp1::hisG, hda1::KanMX, TUP1-3V5::HIS3 |
| FW8430 | MATa, ho::LYS2, lys2, ura3, leu2::hisG, his3::hisG, trp1::hisG, hos1::NatMX, TUP1-3V5::HIS3<br>MATalpha, ho::LYS2, lys2, ura3, leu2::hisG, his3::hisG, trp1::hisG, hos1::NatMX, TUP1-3V5::HIS3 |
| FW8103 | MATa, ho::LYS2, lys2, ura3, leu2::hisG, his3::hisG, trp1::hisG, hos2:HIS3MX, TUP1-3V5::HIS3<br>MATalpha, ho::LYS2, lys2, ura3, leu2::hisG, his3::hisG, trp1::hisG, hos2:HIS3MX, TUP1-3V5::HIS3 |
| FW8457 | MATa, ho::LYS2, lys2, ura3, leu2::hisG, his3::hisG, trp1::hisG, hda1::KanMX, rpd3::HIS3MX, TUP1-3V5::HIS3<br>MATalpha, ho::LYS2, lys2, ura3, leu2::hisG, his3::hisG, trp1::hisG, hda1::KanMX, rpd3::HIS3MX, TUP1-3V5::HIS3 |
| FW8428 | MATa, ho::LYS2, lys2, ura3, leu2::hisG, his3::hisG, trp1::hisG, hos1::NatMX, rpd3::HIS3MX, TUP1-3V5::HIS3<br>MATalpha, ho::LYS2, lys2, ura3, leu2::hisG, his3::hisG, trp1::hisG, hos1::NatMX, rpd3::HIS3MX, TUP1-3V5::HIS3 |
| FW8171 | MATa, ho::LYS2, lys2, ura3, leu2::hisG, his3::hisG, trp1::hisG, rpd3::HIS3MX, hos2:HIS3MX, TUP1-3V5::HIS3<br>MATalpha, ho::LYS2, lys2, ura3, leu2::hisG, his3::hisG, trp1::hisG, rpd3::HIS3MX, hos2:HIS3MX, TUP1-3V5::HIS3 |
| FW4214 | MATa, ho::LYS2, lys2, ura3, leu2::hisG, trp1::hisG, his3::pTEF1-osTIR::HIS3, TUP1-3V5-IAA7::KanMX6, YAP6-3V5::KanMX<br>MATalpha, ho::LYS2, lys2, ura3, leu2::hisG, trp1::hisG, his3::pTEF1-osTIR::HIS3, TUP1-3V5-IAA7::KanMx6, YAP6-3V5::KanMX |
| FW4218 | MATa, ho::LYS2, lys2, ura3, leu2::hisG, trp1::hisG, his3::pTEF1-osTIR::HIS3, TUP1-3V5-IAA7::KanMX6, SOK2-3V5::HIS3<br>MATalpha, ho::LYS2, lys2, ura3, leu2::hisG, trp1::hisG, his3::pTEF1-osTIR::HIS3, TUP1-3V5-IAA7::KanMX6, SOK2-3V5::HIS3 |
| FW5056 | MATa, ho::LYS2, lys2, ura3, leu2::hisG, trp1::hisG, his3::pTEF1-osTIR::HIS3, TUP1-3V5-IAA7::KanMX6, PHD1-3V5::KanMX<br>MATalpha, ho::LYS2, lys2, ura3, leu2::hisG, trp1::hisG, his3::pTEF1-osTIR::HIS3, TUP1-3V5-IAA7::KanMX6, PHD1-3V5::KanMX |
| FW4229 | MATa, ho::LYS2, lys2, ura3, leu2::hisG, trp1::hisG, his3::pTEF1-osTIR::HIS3, TUP1-3V5-IAA7::KanMX6, MOT3-3V5::KanMX<br>MATalpha, ho::LYS2, lys2, ura3, leu2::hisG, trp1::hisG, his3::pTEF1-osTIR::HIS3, TUP1-3V5-IAA7::KanMX6, MOT3-3V5::KanMX |

| | |
|---|---|
| FW4230 | MATa, ho::LYS2, lys2, ura3, leu2::hisG, trp1::hisG, his3::pTEF1-osTIR::HIS3, TUP1-3V5-IAA7::KanMX6, NRG1-3V5::KanMX<br>MATalpha, ho::LYS2, lys2, ura3, leu2::hisG, trp1::hisG, his3::pTEF1-osTIR::HIS3, TUP1-3V5-IAA7::KanMX6, NRG1-3V5::KanMX |
| FW5055 | MATa, ho::LYS2, lys2, ura3, leu2::hisG, trp1::hisG, his3::pTEF1-osTIR::HIS3, TUP1-3V5-IAA7::KanMX6, NRG2-3V5::KanMX<br>MATalpha, ho::LYS2, lys2, ura3, leu2::hisG, trp1::hisG, his3::pTEF1-osTIR::HIS3, TUP1-3V5-IAA7::KanMX6, NRG2-3V5::KanMX |
| FW4224 | MATa, ho::LYS2, lys2, ura3, leu2::hisG, trp1::hisG, his3::pTEF1-osTIR::HIS3, TUP1-3V5-IAA7::KanMX6, SKO1-3V5::KanMX<br>MATalpha, ho::LYS2, lys2, ura3, leu2::hisG, trp1::hisG, his3::pTEF1-osTIR::HIS3, TUP1-3V5-IAA7::KanMX6, SKO1-3V5::KanMX |
| FW3603 | MATa, ho::LYS2, lys2, ura3, leu2::hisG, his3::hisG, trp1::hisG, yap6::NatMX, TUP1-3V5::HIS3<br>MATalpha, ho::LYS2, lys2, ura3, leu2::hisG, his3::hisG, trp1::hisG, yap6::NatMX, TUP1-3V5::HIS3 |
| FW3979 | MATa, ho::LYS2, lys2, ura3, leu2::hisG, his3::hisG, trp1::hisG, sok2::NatMX, TUP1-3V5::HIS3<br>MATalpha, ho::LYS2, lys2, ura3, leu2::hisG, his3::hisG, trp1::hisG, sok2::NATMX, TUP1-3V5::HIS3 |
| FW3991 | MATa, ho::LYS2, lys2, ura3, leu2::hisG, his3::hisG, trp1::hisG, phd1::KanMX, TUP1-3V5::his3<br>MATalpha, ho::LYS2, lys2, ura3, leu2::hisG, his3::hisG, trp1::hisG, phd1::KanMX, TUP1-3V5::his3 |
| FW4239 | MATa, ho::LYS2, lys2, ura3, leu2::hisG, his3::hisG, trp1::hisG, sok2::NatMX, yap6::NatMX, TUP1-3V5::HIS3<br>MATalpha, ho::LYS2, lys2, ura3, leu2::hisG, his3::hisG, trp1::hisG, sok2::NatMX, yap6::NatMX, TUP1-3V5::HIS3 |
| FW4710 | MATa, ho::LYS2, lys2, ura3, leu2::hisG, his3::hisG, trp1::hisG, sok2::NatMX, phd1::KanMX, TUP1-3V5::HIS3<br>MATalpha, ho::LYS2, lys2, ura3, leu2::hisG, his3::hisG, trp1::hisG, sok2::NatMX, phd1::KanMX, TUP1-3V5::HIS3 |
| FW4406 | MATa, ho::LYS2, lys2, ura3, leu2::hisG, his3::hisG, trp1::hisG, phd1::KanMX, yap6::NatMX, TUP1-3V5::HIS3<br>MATalpha, ho::LYS2, lys2, ura3, leu2::hisG, his3::hisG, trp1::hisG, phd1::KanMX, yap6::NatMX, TUP1-3V5::HIS3 |
| FW4010 | MATa, ho::LYS2, lys2, ura3, leu2::hisG, his3::hisG, trp1::hisG, sok2::NATMX, phd1::KanMX, yap6::NATMX, TUP1-3V5::HIS3<br>MATalpha, ho::LYS2, lys2, ura3, leu2::hisG, his3::hisG, trp1::hisG, sok2::NATMX, phd1::KanMX, yap6::NATMX, TUP1-3V5::HIS3 |
| FW5657 | MATa, ho::LYS2, lys2, ura3, leu2::hisG, his3::hisG, trp1::hisG, nrg1::KanMX, sok2::NatMX, phd1::KanMX, yap6::NatMX, TUP1-3V5::HIS3<br>MATalpha, ho::LYS2, lys2, ura3, leu2::hisG, his3::hisG, trp1::hisG, nrg1::KanMX, sok2::NatMX, phd1::KanMX, yap6::NatMX, TUP1-3V5::HIS3 |
| FW5370 | MATa, ho::LYS2, lys2, ura3, leu2::hisG, his3::hisG, trp1::hisG, pIME1-ime1::KanMX, trp1::pNH604-pIME1(WT)-sfGFP-IME1::TRP1, TUP1-3V5::HIS3<br>MATalpha, ho::LYS2, lys2, ura3, leu2::hisG, his3::hisG, trp1::hisG, pIME1-ime1::KanMX, trp1::pNH604-pIME1(WT)-sfGFP-IME1::TRP1, TUP1-3V5::HIS3 |
| FW5372 | MATa, ho::LYS2, lys2, ura3, leu2::hisG, his3::hisG, trp1::hisG, pIME1-ime1::KanMX, trp1::pNH604-pIME1(bsΔ)-sfGFP-IME1::TRP1, TUP1-3V5::HIS3<br>MATalpha, ho::LYS2, lys2, ura3, leu2::hisG, his3::hisG, trp1::hisG, pIME1-ime1::KanMX, trp1::pNH604-pIME1(bsΔ)-sfGFP-IME1::TRP1, TUP1-3V5::HIS3 |
| FW7733 | MATa, ho::LYS2, lys2, ura3, leu2::hisG, his3::hisG, trp1::hisG, pIME1-ime1::KanMX, trp1::pNH604-pIME1(spy3)-sfGFP-IME1::TRP1, TUP1-3V5::HIS3<br>MATalpha, ho::LYS2, lys2, ura3, leu2::hisG, his3::hisG, trp1::hisG, pIME1-ime1::KanMX, trp1::pNH604-pIME1(spy3)-sfGFP-IME1::TRP1, TUP1-3V5::HIS3 |
| FW8079 | MATa, ho::LYS2, lys2, ura3, leu2::hisG, his3::hisG, trp1::hisG, pIME1-ime1::KanMX, trp1::pNH604-pIME1(WT)-sfGFP-IME1::TRP1, YAP6-3V5::KanMX<br>MATalpha, ho::LYS2, lys2, ura3, leu2::hisG, his3::hisG, trp1::hisG, pIME1-ime1::KanMX, trp1::pNH604-pIME1(WT)-sfGFP-IME1::TRP1, YAP6-3V5::KanMX |
| FW8085 | MATa, ho::LYS2, lys2, ura3, leu2::hisG, his3::hisG, trp1::hisG, pIME1-ime1::KanMX, trp1::pNH604-pIME1(bsΔ)-sfGFP-IME1::TRP1, YAP6-3V5::KanMX<br>MATalpha, ho::LYS2, lys2, ura3, leu2::hisG, his3::hisG, trp1::hisG, pIME1-ime1::KanMX, trp1::pNH604-pIME1(bsΔ)-sfGFP-IME1::TRP1, YAP6-3V5::KanMX |
| FW8091 | MATa, ho::LYS2, lys2, ura3, leu2::hisG, his3::hisG, trp1::hisG, pIME1-ime1::KanMX, trp1::pNH604-pIME1(spy3)-sfGFP-IME1::TRP1, YAP6-3V5::KanMX<br>MATalpha, ho::LYS2, lys2, ura3, leu2::hisG, his3::hisG, trp1::hisG, pIME1-ime1::KanMX, trp1::pNH604-pIME1(spy3)-sfGFP-IME1::TRP1, YAP6-3V5::KanMX |
| FW8081 | MATa, ho::LYS2, lys2, ura3, leu2::hisG, his3::hisG, trp1::hisG, pIME1-ime1::KanMX, trp1::pNH604-pIME1(WT)-sfGFP-IME1::TRP1, SOK2-3V5::HIS3<br>MATalpha, ho::LYS2, lys2, ura3, leu2::hisG, his3::hisG, trp1::hisG, pIME1-ime1::KanMX, trp1::pNH604-pIME1(WT)-sfGFP-IME1::TRP1, SOK2-3V5::HIS3 |
| FW8087 | MATa, ho::LYS2, lys2, ura3, leu2::hisG, his3::hisG, trp1::hisG, pIME1-ime1::KanMX, trp1::pNH604-pIME1(bsΔ)-sfGFP-IME1::TRP1, SOK2-3V5::HIS3<br>MATalpha, ho::LYS2, lys2, ura3, leu2::hisG, his3::hisG, trp1::hisG, pIME1-ime1::KanMX, trp1::pNH604-pIME1(bsΔ)-sfGFP-IME1::TRP1, SOK2-3V5::HIS3 |
| FW8093 | MATa, ho::LYS2, lys2, ura3, leu2::hisG, his3::hisG, trp1::hisG, pIME1-ime1::KanMX, trp1::pNH604-pIME1(spy3)-sfGFP-IME1::TRP1, SOK2-3V5::HIS3<br>MATalpha, ho::LYS2, lys2, ura3, leu2::hisG, his3::hisG, trp1::hisG, pIME1-ime1::KanMX, trp1::pNH604-pIME1(spy3)-sfGFP-IME1::TRP1, SOK2-3V5::HIS3 |
| FW8083 | MATa, ho::LYS2, lys2, ura3, leu2::hisG, his3::hisG, trp1::hisG, pIME1-ime1::KanMX, trp1::pNH604-pIME1(WT)-sfGFP-IME1::TRP1, PHD1-3V5::KanMX<br>MATalpha, ho::LYS2, lys2, ura3, leu2::hisG, his3::hisG, trp1::hisG, pIME1-ime1::KanMX, trp1::pNH604-pIME1(WT)-sfGFP-IME1::TRP1, PHD1-3V5::KanMX |
| FW8089 | MATa, ho::LYS2, lys2, ura3, leu2::hisG, his3::hisG, trp1::hisG, pIME1-ime1::KanMX, trp1::pNH604-pIME1(bsΔ)-sfGFP-IME1::TRP1, PHD1-3V5::KanMX<br>MATalpha, ho::LYS2, lys2, ura3, leu2::hisG, his3::hisG, trp1::hisG, pIME1-ime1::KanMX, trp1::pNH604-pIME1(bsΔ)-sfGFP-IME1::TRP1, PHD1-3V5::KanMX |
| FW8095 | MATa, ho::LYS2, lys2, ura3, leu2::hisG, his3::hisG, trp1::hisG, pIME1-ime1::KanMX, trp1::pNH604-pIME1(spy3)-sfGFP-IME1::TRP1, PHD1-3V5::KanMX<br>MATalpha, ho::LYS2, lys2, ura3, leu2::hisG, his3::hisG, trp1::hisG, pIME1-ime1::KanMX, trp1::pNH604-pIME1(spy3)-sfGFP-IME1::TRP1, PHD1-3V5::KanMX |
| FW7650 | MATa, ho::LYS2, lys2, ura3, leu2::hisG, his3::hisG, trp1::hisG, sok2::NatMX, yap6::NatMX, phd1::KanMX, pIME1-ime1::KanMX,<br>trp1::pNH604-pIME1(WT)-sfGFP-IME1::TRP1, TUP1-3V5::HIS3<br>MATalpha, ho::LYS2, lys2, ura3, leu2::hisG, his3::hisG, trp1::hisG, sok2::NatMX, yap6::NatMX, phd1::KanMX, pIME1-ime1::KanMX,<br>trp1::pNH604-pIME1(WT)-sfGFP-IME1::TRP1, TUP1-3V5::HIS3 |
| FW8420 | MATa, ho::LYS2, lys2, ura3, leu2::hisG, his3::hisG, trp1::hisG, sok2::NatMX, yap6::NatMX, phd1::KanMX, pIME1-ime1::KanMX,<br>trp1::pNH604-pIME1(bsΔ)-sfGFP-IME1::TRP1, TUP1-3V5::HIS3<br>MATalpha, ho::LYS2, lys2, ura3, leu2::hisG, his3::hisG, trp1::hisG, sok2::NatMX, yap6::NatMX, phd1::KanMX, pIME1-ime1::KanMX,<br>trp1::pNH604-pIME1(bsΔ)-sfGFP-IME1::TRP1, TUP1-3V5::HIS3 |
| FW8177 | MATa, ho::LYS2, lys2, ura3, leu2::hisG, his3::hisG, trp1::hisG, yap6::NatMX, sok2::NatMX, phd1::KanMX, pIME1-ime1::KanMX,<br>trp1::pNH604-pIME1(spy3)-sfGFP-IME1::TRP1, TUP1-3V5::HIS3<br>MATalpha, ho::LYS2, lys2, ura3, leu2::hisG, his3::hisG, trp1::hisG, yap6::NatMX, sok2::NatMX, phd1::KanMX, pIME1-ime1::KanMX,<br>trp1::pNH604-pIME1(spy3)-sfGFP-IME1::TRP1, TUP1-3V5::HIS3 |
| FW5199 | MATa, ho::LYS2, lys2, ura3, leu2::hisG, his3::hisG, trp1::hisG, pRS306-pCTS1-2xmCherry-SV40NLS<br>MATalpha, ho::LYS2, lys2, ura3, leu2::hisG, his3::hisG, trp1::hisG, pRS306-pCTS1-2xmCherry-SV40NLS |
| FW7473 | MATa, ho::LYS2, lys2, ura3, leu2::hisG, his3::hisG, trp1::hisG, pRS306-pCTS1-2xmCherry-SV40NLS, YAP6-mNeongreen(Yeast Optimized)::NatMX<br>MATalpha, ho::LYS2, lys2, ura3, leu2::hisG, his3::hisG, trp1::hisG, pRS306-pCTS1-2xmCherry-SV40NLS, YAP6-mNeongreen(Yeast Optimized)::NatMX |

| | |
|---|---|
| FW7475 | *MATa, ho::LYS2, lys2, ura3, leu2::hisG, his3::hisG, trp1::hisG, pRS306-pCTS1-2xmCherry-SV40NLS, SOK2-mNeongreen(Yeast Optimized)::NatMX*<br>*MATalpha, ho::LYS2, lys2, ura3, leu2::hisG, his3::hisG, trp1::hisG, pRS306-pCTS1-2xmCherry-SV40NLS, SOK2-mNeongreen(Yeast Optimized)::NatMX* |
| FW7477 | *MATa, ho::LYS2, lys2, ura3, leu2::hisG, his3::hisG, trp1::hisG, pRS306-pCTS1-2xmCherry-SV40NLS, PHD1-mNeongreen(Yeast Optimized)::NatMX*<br>*MATalpha, ho::LYS2, lys2, ura3, leu2::hisG, his3::hisG, trp1::hisG, pRS306-pCTS1-2xmCherry-SV40NLS, PHD1-mNeongreen(Yeast Optimized)::NatMX* |
| FW7644 | *MATa, ho::LYS2, lys2, ura3, leu2::hisG, his3::hisG, trp1::hisG, pRS306-pCTS1-2xmCherry-SV40NLS, TUP1-mNeongreen(Yeast Optimized)::NatMX*<br>*MATalpha, ho::LYS2, lys2, ura3, leu2::hisG, his3::hisG, trp1::hisG, pRS306-pCTS1-2xmCherry-SV40NLS, TUP1-mNeongreen(Yeast Optimized)::NatMX* |
| FW7642 | *MATa, ho::LYS2, lys2, ura3, leu2::hisG, his3::hisG, trp1::hisG, pRS306-pCTS1-2xmCherry-SV40NLS, CYC8-mNeongreen(Yeast Optimized)::NatMX*<br>*MATalpha, ho::LYS2, lys2, ura3, leu2::hisG, his3::hisG, trp1::hisG, pRS306-pCTS1-2xmCherry-SV40NLS, CYC8-mNeongreen(Yeast Optimized)::NatMX* |
| FW5453 | *MATa, ho::LYS2, lys2, ura3, leu2::hisG, his3::hisG, trp1::hisG, tpk1:: tpk1M164G, tpk3::TRP1, tpk2::KanMX6, YAP6-3V5::KanMX*<br>*MATalpha, ho::LYS2, lys2, ura3, leu2::hisG, his3::hisG, trp1::hisG, tpk1:: tpk1M164G, tpk3::TRP1, tpk2::KanMX6, YAP6-3V5::KanMX* |
| FW5454 | *MATa, ho::LYS2, lys2, ura3, leu2::hisG, his3::hisG, trp1::hisG, tpk1:: tpk1M164G, tpk3::TRP1, tpk2::KanMX6, SOK2-3V5::HIS3*<br>*MATalpha, ho::LYS2, lys2, ura3, leu2::hisG, his3::hisG, trp1::hisG, tpk1:: tpk1M164G, tpk3::TRP1, tpk2::KanMX6, SOK2-3V5::HIS3* |
| FW5528 | *MATa, ho::LYS2, lys2, ura3, leu2::hisG, his3::hisG, trp1::hisG, tpk1:: tpk1M164G, tpk3::TRP1, tpk2::KanMX6, PHD1-3V5::KanMX*<br>*MATalpha, ho::LYS2, lys2, ura3, leu2::hisG, his3::hisG, trp1::hisG, tpk1:: tpk1M164G, tpk3::TRP1, tpk2::KanMX6, PHD1-3V5::KanMX* |

**Supplementary Table 2. Oligo nucleotide sequences used througout this study**

| Primer | Oligo sequence (5' to 3') | Targeted region |
|---|---|---|
| *oFW43* | acgatccccgtccaagttatg | *HMR1* (forward) |
| *oFW50* | cttcaaaggagtcttaatttccctg | *HMR1* (reverse) |
| *oFW106* | gtaccaccatgttcccaggtatt | *ACT1* (forward) |
| *oFW268* | agatggaccactttcgtcgt | *ACT1* (reverse) |
| *oFW493* | gatggagggttggcataaaa | 2310 bp upstream of *IME1* (forward) |
| *oFW494* | tgacggtgacgtacgatctcta | 2310 bp upstream of *IME1* (reverse) |
| *oFW248* | ccgtatggtgttggagtaatttg | 2100 bp upstream of *IME1* (forward) |
| *oFW249* | tgccatttagtggacttcttgag | 2100 bp upstream of *IME1* (reverse) |
| *oFW481* | atttttagcgactgccgaaa | 1950 bp upstream of *IME1* (forward) |
| *oFW482* | atgcaacgcctacttgtttt | 1950 bp upstream of *IME1* (reverse) |
| *oFW127* | gccaacttggagaaagaatgtg | 1700 bp upstream of *IME1* (forward) |
| *oFW128* | cggaggtactagtcatcggaat | 1700 bp upstream of *IME1* (reverse) |
| *oFW254* | agaaacgcaaatgctcagagag | 1400 bp upstream of *IME1* (forward) |
| *oFW255* | gaggtaatagcggatgacatcaa | 1400 bp upstream of *IME1* (reverse) |
| *oFW539* | gggtcttaaatacgcagggaat | 1000 bp upstream of *IME1* (forward) |
| *oFW540* | ggcagttcaaaggcttttctta | 1000 bp upstream of *IME1* (reverse) |
| *oFW2685* | aggattgggggtagatacaacatc | 1000 bp upstream of *IME1* (forward) for *pIME1-WT*, *pIME1-bs*, *pIME1-spy* |
| *oFW2688* | gatgggcagttcaaaggct | 1000 bp upstream of *IME1* (reverse) for *pIME1-WT*, *pIME1-bs*, *pIME1-spy* |
| *oFW333* | cttcgagggaaaggatcaaag | 750 bp upstream of *IME1* (forward) |
| *oFW334* | ggctggggggttctgtaattc | 750 bp upstream of *IME1* (reverse) |
| *oFW161* | taaacaacaacaacaacgcaca | 400 bp upstream of *IME1* (forward) |
| *oFW162* | ggcaaggaacaagatcaaaaac | 400 bp upstream of *IME1* (reverse) |
| *oFW463* | caacgcctccgataatgtatatg | *IME1* (forward) |
| *oFW464* | acgtcgaaggcaatttctaatg | *IME1* (reverse) |
